# Successive remodeling of IgG glycans using a solid-phase enzymatic platform

**DOI:** 10.1101/2021.08.27.457991

**Authors:** Yen-Pang Hsu, Deeptak Verma, Shuwen Sun, Caroline McGregor, Ian Mangion, Benjamin F. Mann

## Abstract

The success of glycoprotein-based drugs in various disease treatments has become widespread. Frequently, therapeutic glycoproteins exhibit a heterogeneous array of glycans that are intended to mimic human glycopatterns. While immunogenic responses to biologic drugs are uncommon, enabling exquisite control of glycosylation with minimized microheterogeneity would improve their safety, efficacy and bioavailability. Therefore, close attention has been drawn to the development of glycoengineering strategies to control the glycan structures. With the accumulation of knowledge about the glycan biosynthesis enzymes, enzymatic glycan remodeling provides a potential strategy to construct highly ordered glycans with improved efficiency and biocompatibility. In this study, we quantitatively evaluate more than 30 enzymes for glycoengineering immobilized immunoglobulin G, an impactful glycoprotein class in the pharmaceutical field. We demonstrate successive glycan remodeling in a solid-phase platform, which enabled IgG glycan harmonization into a series of complex-type N-glycoforms with high yield and efficiency while retaining native IgG binding affinity.

**Significance:** Glycosylation plays critical functional and structural roles in protein biology. However, our understanding of how discrete glycan structures affect protein behaviors remains extremely limited due to the naturally occurring microheterogeneity. Through the use of characterized glycoengineering enzyme combination, we report a solid-phase glycan remodeling (SPGR) platform that enables efficient IgG glycan harmonization into several glycoforms of interest with high biocompatibility to the substrates. It provides an efficient strategy to screen the biological behavior of distinct glycoforms, building a fundamental understanding of glycosylation.

## Introduction

Protein glycosylation directly affects the physical and biochemical properties of proteins in eukaryotic systems.(1) According to glycoproteomic analyses, over 1% of the human genome encodes glycosylation-related enzymes and more than 50% of human proteins are glycosylated.(2) Glycoproteins carry structurally diverse oligosaccharides, called glycans, that are involved at the interface of protein-biomolecular interactions and thus determine protein stability, selectivity, and activity. The significance of protein glycosylation to biological systems has been exemplified by several diseases associated with various cancers and the immune system.(3, 4) For example, patients with rheumatoid arthritis were found to have an increased galactosylation level in their serum immunoglobulin G (IgG), though the mechanism remains elusive.(5) Unsurprisingly, it follows that insights into the structure and function of glycans have yielded a profound impact on the development of therapeutic glycoproteins.(6) Manipulating glycan structures present an effective strategy to improve their efficacy and safety by modulating immunological responses, circulatory half-life, and effector functions.(7, 8) Thus, glycoengineering represents a versatile tool and a great opportunity to create better medicines. To achieve this goal, technologies that provide the control of protein glycosylation profile are essential.

However, tools to access the diverse array of glycan structures displayed in nature remain scarce, and methods that provide a high yield of the desired glycoforms have proven to be a still greater challenge to develop despite decades of study.(9, 10) Through traditional synthetic approaches, several common glycoforms have been accessed.(11) These structurally-defined glycans can be installed onto glycoproteins through endoglycosidase and glycosynthase activities.(12, 13) While this approach has advanced our ability to control protein glycosylation, the preparation of synthetic glycans becomes increasingly difficult as the number of saccharide units increases. As a result, the installation of synthetic glycans is not practical for many applications. On the other hand, genetic engineering has been applied for controlled glycan biosynthesis by either knocking out or introducing certain glycoengineering enzymes in the host cells.(14) This strategy enables *in vivo* glycan remodeling and has been demonstrated in non-human cell lines.(15) However, the optimization of this strategy has been impeded by the complexity of engineering glycosylation pathways. Also, micro-heterogeneity is often generated during glycan formation, which, although it is comparable to the natural phenomenon, does not provide exquisite control over the molecular structure (16)

In recent decades, our understanding of the *in vitro* activity of glycoengineering enzymes is growing rapidly.(17–19) Some of the enzymes can even function on intact glycoproteins, which opens a new window for glycan remodeling.(20–22) To further leverage the use of these enzymes, three primary challenges need to be addressed. First, most of the enzymes applied to glycan engineering have been studied based on the use of synthetic oligosaccharides or denatured glycoproteins as substrates.(19, 23) Whether or not they can function on intact glycoproteins needs to be investigated. Second, preserving the integrity and functions of the substrates after the enzymatic reactions is critical, especially for therapeutic glycoproteins. Protocols with high biocompatibility are thus required. Third, to construct complex glycan structures, successive reactions using different enzymes are needed. These enzymes might require very different working conditions, such as pH and temperature. Therefore, one would need to repeat the buffer swapping and product purification processes between the enzymatic reactions, which is highly labor-intensive and time-consuming. Together, to address these issues, novel platforms that enable efficient, successive enzymatic glycan remodeling with high biocompatibility to the substrates are in great demand.

Inspired by solid-phase peptide synthesis (SPPS), herein, we introduce solid-phase glycan remodeling (SPGR) where enzymatic reactions are carried out on the substrates immobilized on resins.(24) This approach enables efficient reaction swapping, substrate purification, and the recovery of both products and engineering enzymes. We use human IgG as the substrate in this study because it is a major class of glycoproteins that have been applied in therapeutic development.(6, 25) We quantitatively examined more than 30 glycan engineering enzymes for their activities on intact IgG immobilized on resins and then applied them in SPGR. This method has allowed us to harmonize IgG glycans into ten different glycoforms, including non-canonical structures, in 48 hours with an average conversion ratio of over 95%. Physical and biochemical analyses indicated that the SPGR-engineered IgGs preserved integrity and functionality, suggesting that SPGR has high biocompatibility to the substrates.

## Results

### The design of solid-phase glycan remodeling (SPGR) for IgG glycoengineering

Our strategy to achieve efficient, successive glycan remodeling is immobilizing IgG onto protein A resins and then executing enzymatic reactions heterogeneously **(Figure 1a)**.^§1^ This enables product purification by filtration, greatly speeding up multi-step reactions. We use empty SPE (solid-phase extraction) columns with standard Luer fittings as the SPGR reaction vessels. The Leur fittings can be connected to either syringes or vacuum manifolds to control the flow speed during washing processes. A frit is inserted into the bottom of the column for trapping the solid supports. SPGR processes can be separated into 5 steps: 1) resin loading; 2) IgG immobilization; 3) washing and conditioning; 4) enzymatic glycan remodeling, and; 5) elution and downstream analyses. The third and fourth steps are repeated for successive reactions until the remodeling is complete.

**Figure 1.**
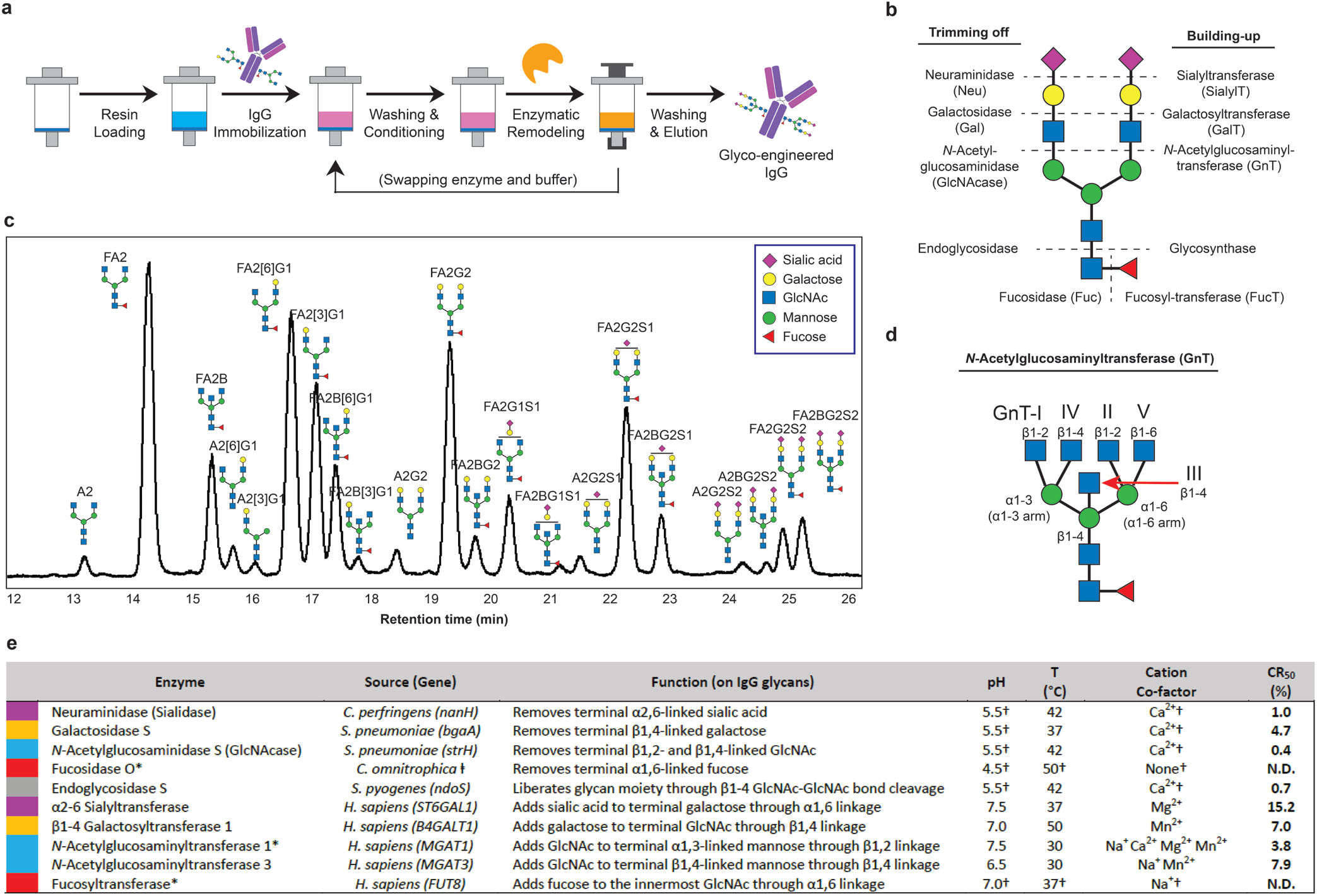
(a) Scheme of solid-phase glycan remodeling (SPGR) protocols. (b) Enzyme map for bi-antennary N-glycan glycoengineering. (c) Chromatogram of glycans collected from human serum IgG. The Oxford notation is used for glycan nomenclature. (d) The functions of *N*-acetylglucosaminyltransferases (GnTs). (e) Glycoengineering enzymes selected for SPGR and their optimized working conditions. CR_50_: the enzyme-to-substrate (IgG) molar ratio required for reaching 50% substrate conversion into the products in one hour using SPGR and the optimized conditions. Enzymes are color coded to indicate the saccharide residues they build or trim. Please refer to **Table S2** for detailed working conditions. N.D.= no data. † The conditions suggested by the enzyme suppliers were used without further optimization. * Glycoengineered IgG was used as the substrate for the characterization. ‡ Genbank KXK31601.1

To identify capable glycoengineering enzymes for SPGR, we quantitatively analyzed the activity of 34 candidates, including exoglycosidases, endoglycosidases, and glycosyltransferases **(Figure 1b,e)**. Each enzyme was incubated with immobilized IgG for 1 or 24 hours. The enzyme activity—indicated by the consumption of substrate glycan species—was then quantified via chromatographic analysis. The candidates with the highest activity in each enzyme class were selected for SPGR applications and their working conditions were further optimized **(Figure 1e, Figure S1-11)**. Also, we defined CR_50_ to be the enzyme-to-substrate (IgG) ratio that leads to 50% substrate conversion into the products in one hour using SPGR **(Figure 1e)**. This value allows us to estimate how much enzyme is required for SPGR reactions when the amount of substrate varies. It also provides the information about the reaction efficiency between different enzymes: the smaller the CR_50_ value, the more efficient the reaction is.

### Trimming IgG glycans with glycosidases

IgGs have two highly conserved glycosylation sites on the crystallizable region (Fc) at Asn 297 where more than 20 complex-type glycoforms have been found with the majority in bi-antennary structures **(Figure 1c)**.(26) IgG glycans play essential roles in Fc receptor (FcR)-mediated activities, such as antibody-dependent cellular cytotoxicity (ADCC).(27) About 20% of IgG glycans contain terminal sialic acids through α2-6 linkages. These sialylated glycans have been known to confer anti-inflammatory activity.(28) Similarly, IgG galactosylation modulates inflammatory properties and about 70% of the IgG glycans contain terminal β1-4 galactoses.(3, 5) The galactosylation level is also known to influence the clearance rate of glycoproteins in serum, mediated by asialoglycoprotein receptors, resulting in a direct impact on their pharmacokinetic properties.(29, 30) Compared to sialic acid and galactose, our understanding of *N*-Acetylglucosamine (GlcNAc)’s impacts on IgG is more limited. GlcNAc exists in all N-glycans and plays decisive roles in glycan biosynthesis pathways. Extended from the chitobiose core, GlcNAc glycosidic linkage serves as a watershed that determines the subclasses of N-glycans: complex-type, high-mannose, and hybrid-type N-glycans. Complex-type glycans can further branch into bisecting, bi-antennary, tri-antennary, and tetra-antennary glycans, and so on.(31) About 90% of the IgG glycans are bi-antennary while the rest of them have bisecting structures.(26)

To control IgG glycoforms, we first aimed to harmonize them into the core saccharides by removing terminal sialic acid, galactose, and then GlcNAc.^§2^ Neuraminidase (Neu, or sialidase) is a class of enzyme that cleaves the glycosidic linkages of sialic acids. Our screening showed that Neu from *Clostridium perfringens* has the highest activity on immobilized IgG with a CR_50_ of 1%. This enzyme has a broad substrate spectrum and can function on all the IgG glycoforms containing terminal sialic acid **(Figure 2, S1)**. Next, galactosidase (Gal) from *Streptococcus pneumoniae* showed the highest activity in our screening (CR_50_= 4.7%) for removing galactose **(Figure S2)**. It functions on all the IgG glycoforms containing terminal galactoses with an optimal temperature at 37°C. To trim off GlcNAc, *N*-Acetylglucosaminidase (GlcNAcase) from *S. pneumoniae* showed the highest activity (CR_50_ 0.4%, **Figure S3**). It has low glycosidic linkage selectivity and can trim terminal GlcNAc extended from the chitobiose core **(Figure 2)**. Sequential treatments using these three enzymes leads to IgG glycan harmonization into (F)M3 structures **(Figure 3a)**.

**Figure 2.**
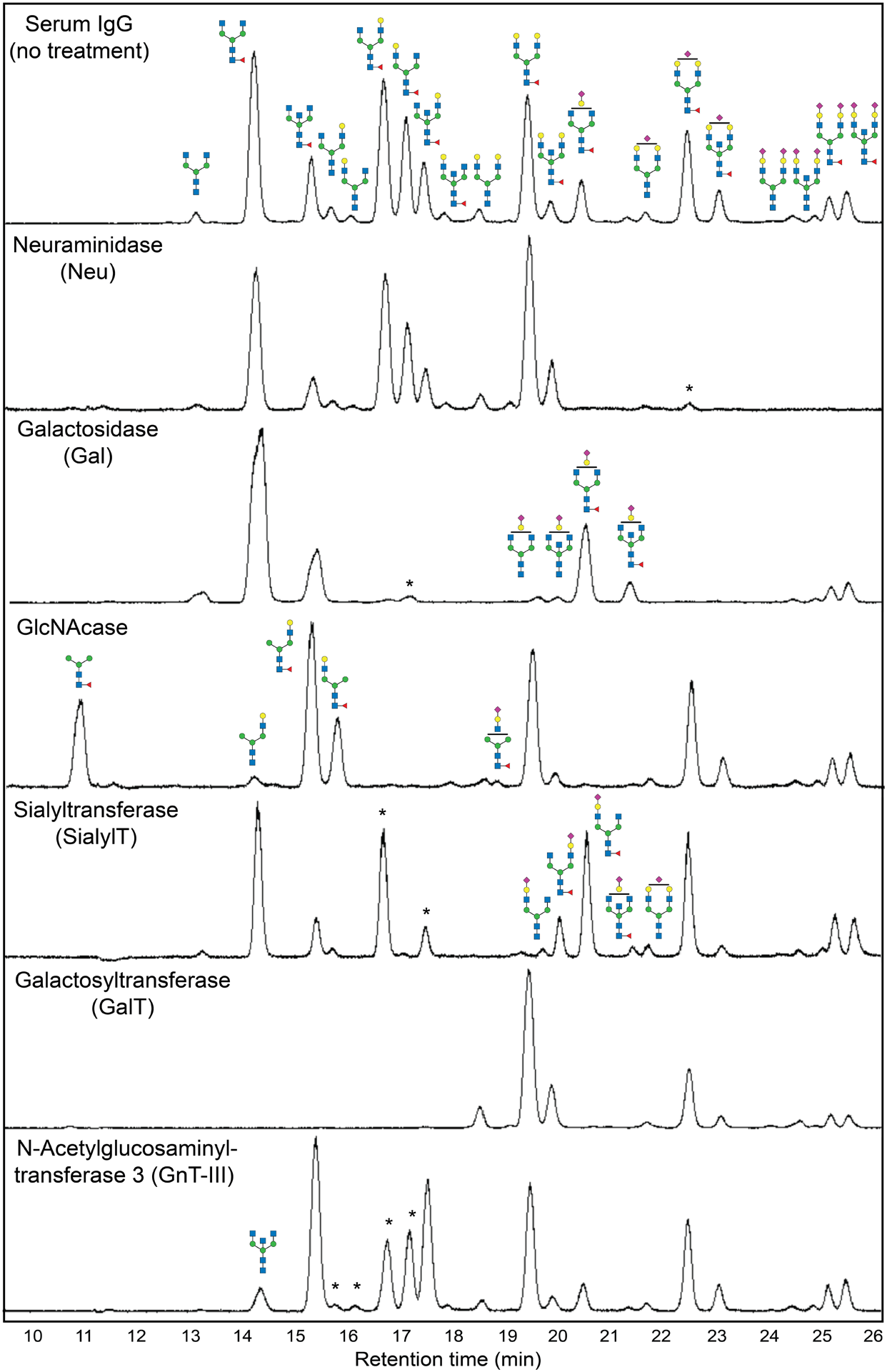
Chromatogram of glycans collected from IgG treated with different glycoengineering enzymes. The data was collected from reactions that reached, or were close to, the plateau of the conversion. The formation of glycoforms was confirmed by mass spectrometry analyses. Please refer to **Table S2** for detailed reaction conditions. Star marks indicated the substrate glycan species that have not been fully transformed in the reaction.

**Figure 3.**
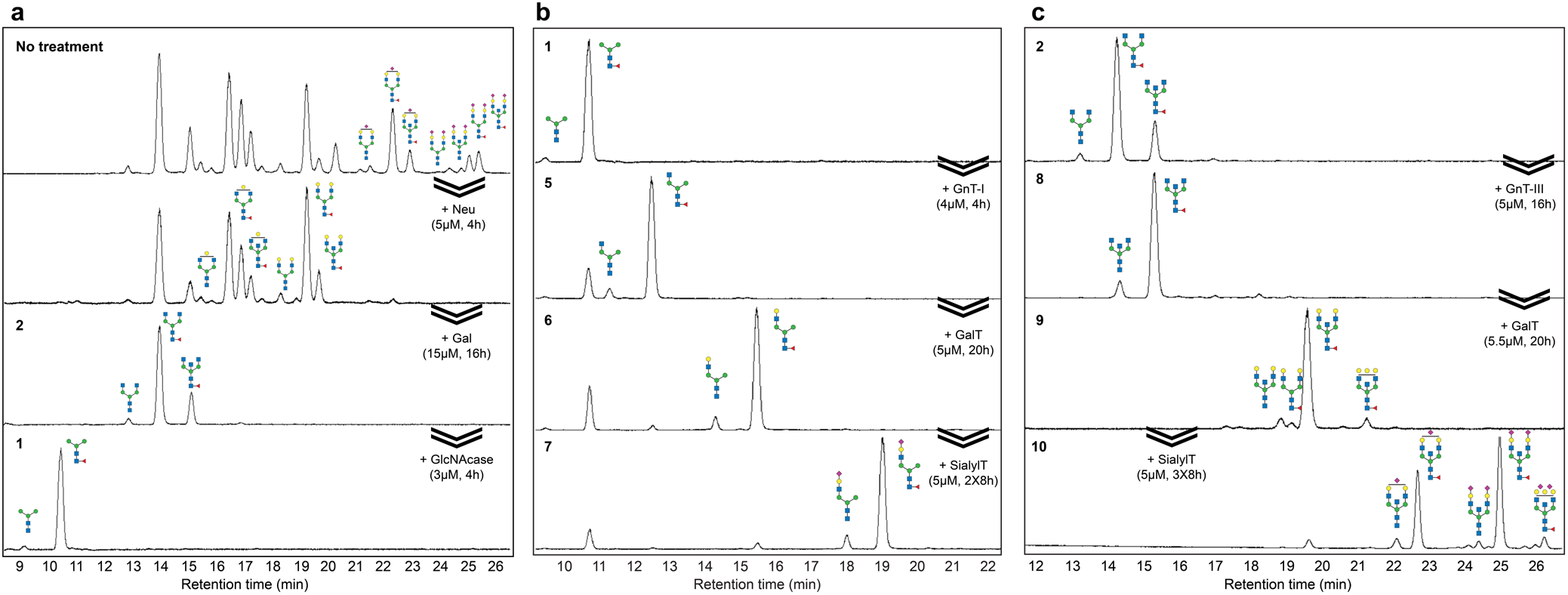
Glycan chromatograms from IgGs after sequencial glycoengineering using SPGR. A) Process of remodeling IgG glycans into core saccharides (FM3 and M3 glycoforms). B) Process of re-building core saccharides into mono-antennary species. About 10% (F)M3 glycans remained in the products due to the reversible activity of GnT-I. C) Process of re-building FA2 and A2 glycans into bisecting species. Refer to **Figure 4** for the sample numbering and **Figure 1** for the buffer conditions.

Fucose on IgG glycan chitobiose core has been known to modulate IgG binding affinity to Fc receptors.(32) Defucosylated IgG has been reported to have an over 50-fold increase in ADCC activity.(33) As such a strong regulator, controlling the level of IgG core fucose has become an attractive strategy for improving the efficacy of IgG-based drugs. Over 90% of the human serum IgG glycans are fucosylated.(26) To identify the enzymes that can trim fucose from intact IgGs in their native confirmations, we tested seven fucosidases (Fuc). Unfortunately, none of them showed an acceptable activity **(Table S1)**. It has been reported that Fuc only functions on intact IgG when their glycans are trimmed down to the GlcNAc-fucose disaccharides, which indicates a strong steric interference between the Fuc-fucose interaction.(20) Inspired by works from *Huang et al*., we tested the Fuc panel with glycoengineered IgG bearing (F)M3 glycans, as prepared above. The enzyme from *Candidatus omnitrophica* showed significantly improved activity on this group of substrates **(Figure S4)**. ^§3^ A 20% conversion was achieved in a 3-day reaction. The conversion ratio was further increased to 65% if non-immobilized substrates were used.

### Building IgG glycans with glycosyltransferases

Glycosyltransferases catalyze the transfer of saccharide(s) from activated sugar phosphates, the glycosyl donors, to glycosyl acceptor molecules, such as glycoproteins.(34) Sialyltransferase (SialylT) from *Homo sapiens* exhibited the highest activity in our screening for installing sialic acid through α2-6 linkage to the IgG with terminal galactose. This enzyme has a relatively high CR_50_ of 15.2% with an apparent substrate selectivity, as shown in **Figure 2** and **Figure S5**. Di-galactosylated glycan (FA2G2) and mono-galactosylated glycan with galactose at the α1-3 arm (FA2[3]G1) were completely transformed after a 16-hours reaction; while mono-galactosylated glycans at the α1-6 arm (FA2[6]G1) showed only minimal sialylation. The selective sialylation observed here agreed with previous reports and was likely caused by the folded conformation that the Fc region adopts when the galactose on the α1-6 arm is present.(35, 36) Besides, we also observed a decreased enzyme activity when the (F)A2G2 glycans were mono-sialylated **(Figure S5)**.

To install galactose on IgG glycans, we selected the galactosyltransferase (GalT) from *Homo sapiens* **(Figure 2, S6)**.(37) This enzyme catalyzed the transfer of galactose from Uridine 5’-diphosphogalactose (UDP-Gal) to IgG glycans with terminal GlcNAc. It has a CR_50_ of 7% and a broad spectrum of substrate specificity that enables the transformation of all the non- and mono-galactosylated IgG glycans into bi-galactosylated forms.

The addition of GlcNAc to the chitobiose core is relatively complicated because this process involves a series of *N*-Acetylglucosaminyltransferases (GnT) with various substrate specificities **(Figure 1d)**.(38) We investigated the activity of five human GnTs that are responsible for complex N-glycan synthesis and obtained positive results from GnT-I, III, and V. GnT-I (MGAT1) initiates the formation of the complex-type and hybrid N-linked glycans by installing an β1-2 GlcNAc to the α1-3 mannose **(Figure S7)**.(39) We observed a good CR_50_ of 3.8% in our activity screening, calculated based on (F)M3 glycan consumption. This enzyme likely possesses GlcNAcase activity as well because the conversion ratio reached a plateau of ~85% **(Figure S8)** in all the conditions we tested. Whether or not its GlcNAcase activity can be repressed through genetic engineering to reach full conversion remains to be investigated. Human GnT-III (MGAT3) serves to install the bisecting GlcNAc to the β1-4 mannose through a β1-4 linkage.(40) A higher level of bisecting GlcNAc on IgG results in enhanced ADCC activity and immune cells effector functions.(41) Reactions using serum IgG as the substrate suggested that human GnT-III can function on IgG glycoforms containing at least one terminal GlcNAc **(Figure 2)**. Moreover, it showed much higher activity on glycans with two terminal GlcNAcs as opposed to a single GlcNAc **(Figure S9)**.

Tri- and tetra-antennary N-glycans are not typically reported on native human serum IgG, and were not observed in our studies. Human GnT-V (MGAT5) is reported to add the secondary GlcNAc to the α1-6 mannose through β1-6 linkage and leads to the formation of tri-/tetra-antennary glycoforms.(42) We observed GnT-V activity after a 24-hours reaction with intact IgG, revealed by the formation of tri-antennary species. **(Figure S10)**. The activity of GnT-V on intact IgG is low but potentially can be improved through genetic control or evolution for future applications.

Finally, mammalian α1,6-fucosyltransferase (FucT) catalyzes the transfer of a fucose residue from GDP-fucose, the donor substrate, to the reducing-end terminal GlcNAc residue through an α1,6-linkage.(43) The activity of FucT on intact IgG was observed but the conversion ratio was low, likely due to the strong steric hindrance on the substrate. We further boosted the FucT reaction by increasing both enzyme concentration and incubation time. The result indicated that FucT prefers IgGs bearing A2 and A2G1 glycoforms **(Figure S11)**. Our data suggested that, by employing Fuc and FucT, modulating the level of core fucose on intact IgG is feasible.

### Reconstructing a harmonized glycosylation profile

It was described how a Neu reaction followed by Gal treatment turned IgG glycans into GlcNAc-terminating glycoforms **(Figure 3a)**. We can further run a GlcNAcase reaction to trim the glycans into the core structures terminating with mannose. The whole sequence was performed in 24 hours without isolating IgG between steps. Similarly, remodeling using Neu and then GalT generated glycans with terminal galactose **(Figure S12a)**; while a GalT reaction followed by SialylT resulted in mono- and bi-sialylated species **(Figure S12b)**. The demonstrated SPGR routes in this work are summarized in **Figure 4**.

**Figure 4.**
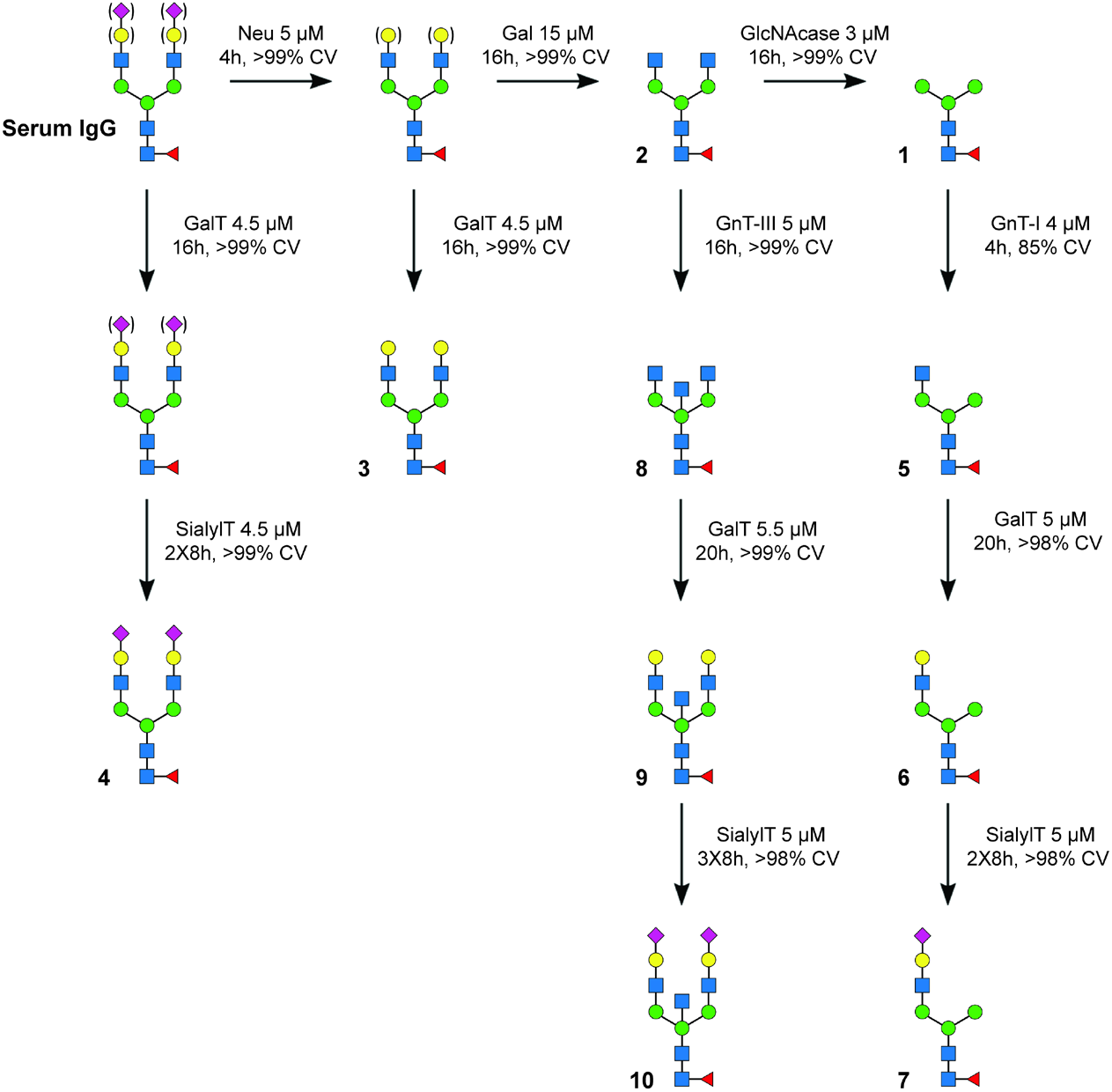
Scheme of SPGR routes investigated in this work. Bracket: with or without the residues (microheterogeneity). Serum IgG contains a ~5% defucosylated population and a ~10% bisecting population. For simplicity, these two populations are not shown in the scheme. The conversion (CV) ratio was calculated based on the consumption of substrate glycan species.

IgG bearing (F)M3 glycans can also serve as starting materials for rebuilding non-canonical glycoforms. Starting with the (F)M3 glycans, we applied GnT-I, GalT, and then SialylT reactions to construct a series of mono-antennary species **(Figure 3b)**. Mono-antennary glycans are rare in nature and their effects on protein biology remain elusive. Whether they can be utilized for regulating the interactions between IgG and FcR is a great research topic of interest. In addition to mono-antennary species, we also thought to increase the population of bisecting glycans on IgG. We trimmed off terminal sialic acids and galactose from IgG and then introduced GnT-III to the resulting (F)A2 glycans. After overnight incubation at 5 μM, we reached a full conversion of IgG glycans into the bisecting forms **(Figure 3c)**. These bisecting glycans can also be further galactosylated or sialylated using SPGR. Compared to mono-antennary species, both GalT and SialylT showed slightly decreased activity on bisecting glycans. Therefore, a higher enzyme concentration or longer incubation time is required to achieve full conversion. The preparation of synthetic glycans with complex structures, traditionally, not only requires extensive experience but also a lot of time and effort. SPGR allowed us to reconstruct IgG glycans into the mono-antennary and bisecting glycoforms within 2-3 days, presenting a very efficient strategy for glycoengineering.

### SPGR is biocompatible

To examine the biocompatibility of SPGR to the substrates, we analyzed the physical properties of the glycoengineered IgGs prepared above. Size analyses using dynamic light scattering (DLS) showed no significant difference between native serum IgG and SPGR-engineered IgGs, suggesting that there was no denaturing and/or aggregation occurred during the remodeling processes **(Figure 5a)**. On the other hand, slight changes in their melting temperature (T_m_) and aggregation temperature (T_agg_) were observed **(Figure 5b-c, Table S3)**. We reasoned these changes were attributed to the altered structure of IgG. It has been known that the Asn297 glycans play roles in maintaining the conformation and stability of the Fc region through intra-molecular interactions.(44, 45) Decreased T_m_ and T_agg_ values were observed in IgGs terminating with mannose **(1)** or GlcNAc **(2,5,8)**, the glycoforms showing reduced intra-molecular interactions.(44) Similarly, our mono-antennary glycans **(5-7)** had lower T_m_ values compared to others. These glycoforms lack the α1-6 arm which is important for forming intra-molecular interaction within the Fc region.(35) As expected, the removal of the bulk of the glycan structures using Endo S led to a dramatic drop in T_m_ and T_agg_.

**Figure 5.**
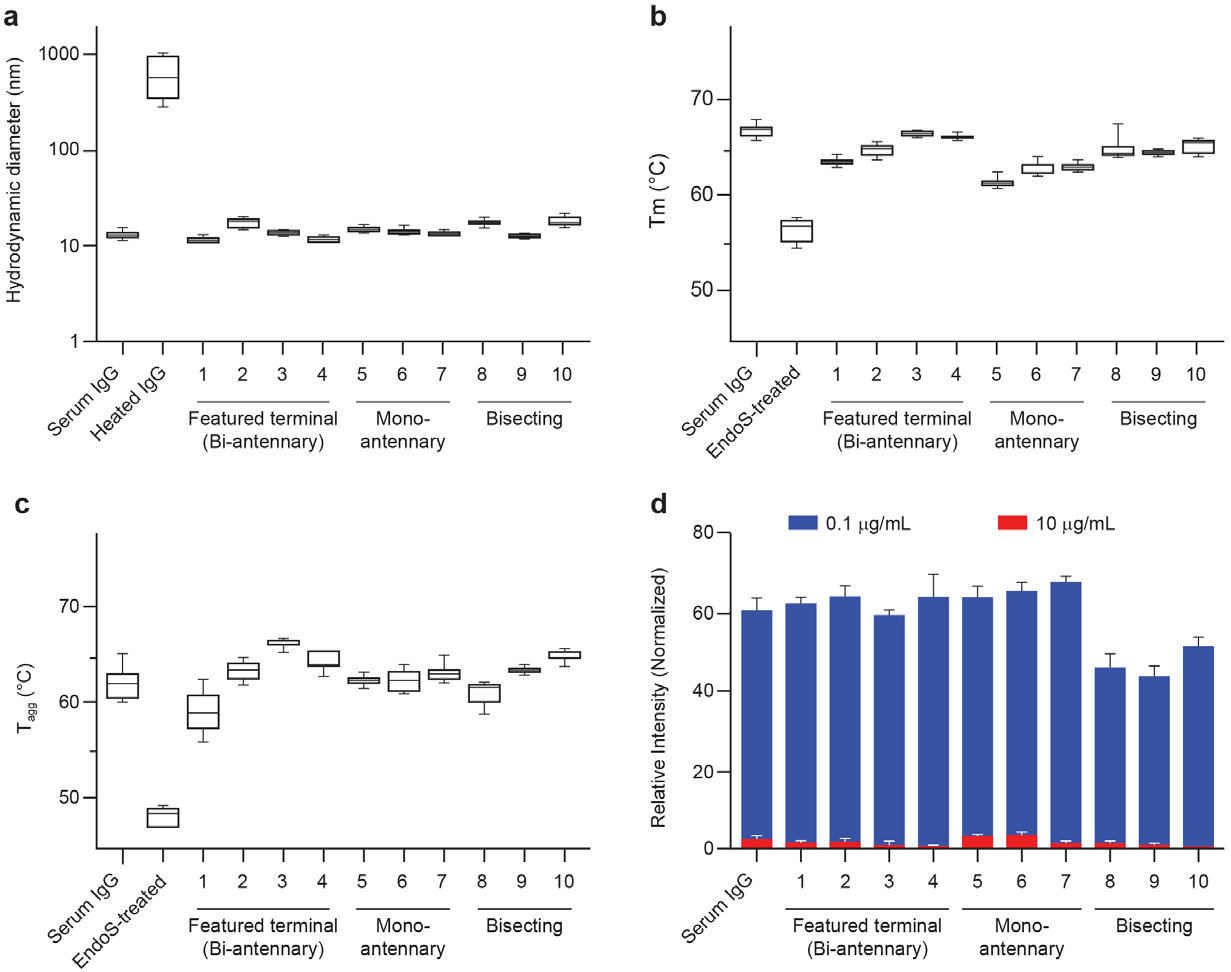
Physical and biochemical characterization of SPGR-engineered IgGs. (A) Size analyses using dynamic light scattering (DLS). (B) Melting temperature analyses (T_m_). (C) Aggregation temperature analyses (T_agg_). Endo S treatment removed most IgG glycan structures, leaving only GlcNAc-Fuc di-saccharide on IgG. (D) Competition assay revealed the binding affinity of SPGR-engineered IgG to Fc gamma receptor I (FcγR I). The lower the relative intensity, the stronger the interaction between SPGR-engineered IgG and FcγR I was. Refer to **Figure 3** and **S12** for the chromatograms of analyzed samples.

Next, we tested whether SPGR-engineered IgGs preserve the binding ability to FcRs. We performed a competition assay where the interaction between SPGR-engineered IgGs and Fc gamma receptor I (FcγR I, or CD64) resulted in fluorescence signal reduction. A signal reduction of 60% was found in all the glycoengineered IgGs at the concentration of 0.1 μg/ml; while complete inhibitions were reached at about 10 μg/ml **(Figure 5D, Figure S13)**. Furthermore, the bisecting glycoforms **(8-10)** showed decreased EC_50_ values in this assay, indicating an enhanced binding affinity to FcγR I **(Table S3)**. Previous studies have reported that increased level of bisecting glycoforms in mouse IgG1 results in enhanced ADCC activity, possibly owing to improved FcγR III binding.(41) How the bisecting GlcNAc enhances FcγR I and III bindings, and whether or not through the same mechanisms, remain elusive but intriguing. Together, our characterization analyses suggested that: 1) SPGR-engineered IgGs have preserved integrity and functions, and 2) the difference in FcR binding affinity between mono-antennary, bi-antennary, and bisecting glycoforms could present a useful handle to control IgGs’ immunogenicity.

## Discussions

For decades there has been a clear demand for glycoengineering tools that enable scholars to explore the role(s) of glycan structures on the function and form of glycoconjugates. SPGR presents a straightforward strategy for controlling glycan structures with several advantages: 1) it is a biocompatible approach with minimal disruption to the protein substrates; 2) it circumvents the need to prepare synthetic glycans, which can be cumbersome; 3) tight control of glycoforms is achievable with the use of different enzyme combinations and; 4) the procedures are user-friendly and can be readily automated, greatly reducing the cost for industrial applications. Moreover, the idea of executing sequential enzymatic remodeling on immobilized proteins can potentially be extended to most existing biocatalytic cascade reactions involving different classes of enzymes and substrates.(46)

Beyond the presented work, there remains an opportunity to explore alternative immobilization strategies in future experiments. IgG immobilization using protein A, a 47kD protein, likely limited the enzyme efficiency by creating a strong steric hindrance.^§4^ Methods using oligopeptides, such as a his-tag, presumably have a lower steric effect and can enable higher enzyme activities. Since immobilization has been commonly employed for protein purification in pharmaceutical manufacturing processes, SPGR can conceivably be inserted into modern protein production processes as a “glycan modification module” to provide pure, glycoengineered proteins.

A particularly disruptive application of SPGR would be to humanize glycoforms on therapeutic proteins produced from non-human cell lines. Chinese hamster ovary (CHO) is commonly used for therapeutic protein production because they generate human-like post-translational modifications.(47) However, non-human glycoforms still exist in the cell line and should be removed to reduce the potential immune response in patients.(48) SPGR can be employed to humanize those glycans during the production process. From a different perspective, protein production in mammalian hosts is costly because of its long fermentation time and liability of virus infections. To address this issue, yeast has been employed as an alternative host for large-scale expression of therapeutic proteins. Glycoproteins expressed from yeast contain high-mannose N-glycans which confer a short half-life *in vivo* and thereby compromise the efficacy of most therapeutic proteins.(7) Therefore, gene engineered strains are constructed for producing human-mimicking glycan patterns.(49) To date, a couple of simple glycoforms have been achieved in yeast and they have provided the desired scaffold (e.g. M5 glycan) for downstream glycan remodeling *in vitro*.(15) With the combination of SPGR, large-scale production of therapeutic proteins in yeast with controlled glycan structures is possible.

The awareness of public health has been raised significantly in the past months, intensifying the demand for developing better biologic medicine. Glycoproteins have proven their unignorable values in therapeutic and vaccine development, which stresses the urgent need for exquisite control of glycosylation profile for improved safety and efficacy. SPGR presents an efficient, user-friendly method for glycoengineering. It enables the control of glycan structures with various glycoforms and, presumably, on diverse glycoproteins. We believe that SPGR will greatly accelerate protein glycosylation studies as well as their pharmaceutical applications.

## Data Availability

All data are available upon reasonable request to the corresponding authors.

## Author contributions

SPGR method development, enzyme activity screening/characterization, and data analyses were performed by Y-P. H. Computational modeling was done by D.V. All the authors were involved in the design of the research and manuscript drafting.

## Acknowledgment

We thank the Merck Research Laboratory (MRL), Merck Postdoctoral Research Fellows Program, and Analytical Research and Development (AR&D) department for the financial support. We are also thankful to Anumita Saha (Data-Rich Measurement group), Alycia Shoultz (Protein Engineering group), Jacob Forstater (Chemistry Enabling Technology group), and Fuh-Rong Tsay (Method Screening and Purification group) for technical assistance and advice.

## Footnotes

§ See supplementary information for more discussions.

