## Supplementary materials for "Successive remodeling of IgG glycans using a solid-phase enzymatic platform"

Raw data that support the findings of this study are available upon reasonable request.

**SI-Materials and Instruments**

Human serum IgG (I4506), Tris-HCl (T-5941), HEPES (H4034), Sodium acetate (S2889), Calcium chloride (C5670), Magnesium chloride (M4880), Manganese Chloride solution (M1787), Sodium chloride solution (S5150), CMP-NANA (C8271), UDP-Gal (U4500), UDP-GlcNAc (U4375), GDP-Fuc (G4401), Acetonitrile (900667), Discovery glycan SPE columns (55465-U), and empty SPE column and frit (57607-U) are purchased from Sigma-Aldrich. Protein A resin (53139), Protein A-IgG binding buffer (54200), Protein A-IgG elution buffer (21027), $H_2O$ (10977015) were purchased from ThermoFisher. HILIC columns for chromatography (186004742), SPE µPlate (186002780), µPlate extraction manifold (186001831), SPE vacuum manifold (WAT200607), RapiGest SF (186008090), Glycoworks buffer (186008100), glycan quantitative standard (186008791), and Rapifluor-MS (186008091) were purchased from Waters. MWCO filters (UFC503096) were purchased from Millipore Sigma. Rapid PNGaseF (P0710) was purchased from NEB. Human FcR AlphaLISA Binding Kit (AL3081C) was purchased from PerkinElmer. Please refer to **Table S1** for the vendor information of glycoengineering enzymes. No unexpected or unusually high safety hazards were encountered in this work.

Chromatography and mass spectrometry analyses were conducted using an Agilent 1290 Infinity II LC system tandem with Agilent 6500 Series quadrupole time-of-flight MS system. Enzyme concentration was determined by absorbance at 280 nm using NanoDrop 2000 (Thermo Scientific). Temperature-controlled reactions/incubations were performed in ThermoFisher MaxQ 6000 incubator and Fisherbrand Thermal Mizer II. Measurements of hydrodynamic diameter, melting temperature ($T_m$) and aggregation temperature ($T_{agg}$) were performed on Uncle (all-in-one biologics stability screening platform) from Unchained Labs. The detection of AlphaLISA-based assays for IgG-FcγR binding studies was conducted by EnVision Multimode Plate Reader 2105 (PerkinElmer).

**SI-Methods**

Solid-phase glycan remodeling (SPGR)

**(I) Resin loading:** Empty SPE columns composed of an empty column body, a frit with 0.2 µm pores, and a lid were used as reaction vessels for SPGR reactions. The columns were mounted onto a 20-wells SPE vacuum manifold. 100 µl of protein A resin (wet resin) was transferred into each column, followed by conditioning with 0.8 ml protein A-IgG binding buffer twice. A vacuum system was connected to the manifold to control the flow rate. **(II) IgG immobilization:** 1 mg (unless other specified) human serum IgG was added to 0.5 ml protein A-IgG binding buffer, followed by gently shaking until all the powder was dissolved. The solution was then transferred to the SPE column containing protein A resins. To ensure good immobilization, the columns were dismounted from the manifold, capped with Luer fittings, and then incubated for 15 minutes at room temperature with gentle rotating. **(III) Washing and conditioning:** After the incubation, the columns were mounted to the manifold again, followed by washing with 0.8 ml protein A-IgG binding buffer (3 times) and then enzyme reaction buffer (2 times). After the conditioning step, the buffer was completely drained out from the columns. **(IV) Enzyme reactions:** Enzyme reaction solutions were prepared by mixing the desired amount of enzyme, reaction buffer, and saccharide donors (1 mM, for glycosyltransferase) to a final volume of 100 µl unless other specified. Please refer to **Figure 1e** and **Table S2** for the details of reaction conditions, including pH, concentration, and cation cofactors. Enzyme reactions were initiated by transferring the reaction solution into SPE columns that contain immobilized IgG substrates. The columns were capped by Luer fitting, sealed with parafilm, and incubated at temperature-controlled shakers for a certain amount of time **(Table S2)**. After the reaction, the enzyme solution was discarded (or recovered) and the columns were washed 6-8 times with 0.8 ml protein A-IgG binding buffer via the extraction manifold. **(V) Elution of glycoengineered IgG:** 500 µl protein A-IgG elution buffer was added to the reaction columns in two portions with a 2-minutes incubation for each at room temperature. The eluent was collected into a 0.5 ml MWCO (molecular weight cut-off) tube and concentrated via centrifugation at 14000g for 5 minutes, followed by buffer exchange into 50 mM HEPES buffer (pH 8). After adjusting the concentration of the eluted IgG substrate to 3 mg/ml using NanoDrop, the IgG substrates were stored at 4°C and were ready for analysis. IgG concentration was adjusted in this step in order to ensure the same amount of sample was charged to the downstream analyses.

LC-MS Analysis of IgG glycans

This protocol is adapted from the Glycoworks manual provided by Waters. **(I) Glycan isolation:** 7.5 µl IgG substrate (3 mg/ml, prepared as described above), 6 µl RapiGest SF (50 mg/ml, in Glycoworks buffer), and 15.3 µl water were mixed in a 1.5 ml microtube. The mixture was then incubated at 90 °C for 5 minutes to denature the substrates. After the samples were cooled down to room temperature, 1.2 µl Rapid PNGase F was added to the tube, followed by another incubation at 50 °C for 10 minutes. **(II) Glycan labeling:** After PNGase F digestion, 12 µl RapiFluor-MS solution (70 mg/ml, in DMF) was added to the solution. The mixture was gently vortexed and then incubated at room temperature for 20 minutes without any light exposure. **(III) Glycan purification:** After labeling, the samples were diluted with 360 µl acetonitrile (1:9 volume ratio). Oasis SPE µPlate from Waters (along with the use of µPlate extraction manifold) was employed for the 1st solid-phase extraction purification, and Discovery SPE from Sigma-Aldrich (along with the use of 20-wells SPE vacuum manifold) was used for the 2nd purification to ensure high signal-to-noise ratio: The SPE columns/µPlate were first washed by water (1 column volume) and then conditioned by water-acetonitrile solution (10:90 v/v, 1 column volume). The glycan samples (in 90% acetonitrile solution) were then charged to the column/µPlate, followed by washing with washing buffer (formic acid/water/acetonitrile 1:9:90 v/v/v, 2 column volume). The glycans were then eluted using 60 µl elution buffer (200 mM ammonium acetate in 5% acetonitrile). **(IV) HILIC-MS analysis:** The purified glycan samples were injected to UPLC equipped with ACQUITY BEH Glycan column (130 Å, 1.7 µm, 2.1 x 150 mm) tandem with IMQ-TOF MS for glycan profile analysis. Please refer to the literature published by Pucic *et. al.* and Kristic *et. al.* for detailed peak assignment.(1, 2) The method provided by Waters for glycan chromatography was used in this work:

**Mobile phase A:** 50 mM ammonium formate in $H_2O$, pH 4.4
**Mobile phase B:** 100% Acetonitrile
**Temperature:** 60 °C
**Injection volume:** 5-10 µl

| Time (min) | Flow rate (ml/min) | %A | %B |
|---|---|---|---|
| 0 | 0.4 | 25 | 75 |
| 35 | 0.4 | 46 | 54 |
| 36.5 | 0.2 | 100 | 0 |
| 39.5 | 0.2 | 100 | 0 |
| 43.1 | 0.2 | 25 | 75 |
| 47.6 | 0.4 | 25 | 75 |
| 55 | 0.4 | 25 | 75 |

Conversion ratio quantification

The conversion ratio of each SPGR reaction was calculated based on the consumption of the substrate glycan species. UV absorption at 260 nm (RapiFluor-MS) from chromatography analysis was used for the quantification of glycan populations. We first normalized the chromatographic peak area of the substrate glycan species to the total glycan peak area (Equation 1). This gives us the percentage of substrate glycan population. The reduction of the substrate glycan population after the reaction was then divided by the initial value to calculate the percentage of substrate conversion (Equation 2). We assume that there is no glycan shedding off from IgG during the experiments (namely, no endoglycosidase activity). Please refer to **Table S4** for detailed substrate species used in the calculation.

For endoglycosidase reactions, their activity was quantified by absolute glycan quantification using an internal standard. This is because endoglycosidases' activity results in the reduction of all glycan signals in the chromatography analyses. A known amount of internal standard (Glycan quantitative standard, Waters) was added to the sample to measure the amount of IgG glycan before and after the reactions.

$$\text{Substrate glycan population} = \frac{\sum \text{Peak area}_{\text{substrate } 1,2,3\ldots i}}{\sum \text{Peak area}_{\text{all glycans}}} = \text{R}_\text{s} \qquad \text{(Equation 1)}$$

$$\text{Conversion Ratio} = \frac{\text{R}_{\text{S (initial)}} - \text{R}_{\text{S (final)}}}{\text{R}_{\text{s (initial)}}} \qquad \text{(Equation 2)}$$

<u>$CR_{50}$ calculation</u>

$CR_{50}$ is defined as the enzyme-to-substrate (IgG) molar ratio that leads to 50% substrate species conversion into the products in 1 hour at optimized working conditions (temperature, pH, cation) using SPGR. This value is determined based on dose-dependent experiments where the conversion ratio at different enzyme concentrations in 1 hour was tested. A sigmoidal curve fitting ([agonist] vs normalized response, GraphPad Prism, see the equation below) was then applied to the data for calculating the enzyme concentration that gives 50% substrate conversion. The resulting enzyme concentration is divided by the substrate (IgG) concentration to give $CR_{50}$.

$$\text{Y} = \frac{100\text{X}}{\text{CR}_{50} + \text{X}}$$

, where Y is normalized response from 0 to 100; X is the concentration of enzyme.

<u>Physical property characterization of SPGR-engineered IgGs</u>

Dynamic light scattering (DLS), melting temperature ($T_m$) and aggregation temperature ($T_{agg}$) studies were executed using UNcle (Unchainedlabs). Glycoengineered IgGs from SPGR reactions were eluted from protein A resins, followed by buffer exchange into HEPES buffer as described above. The protein concentration was then adjusted to 1 mg/ml using NanoDrop. 9 µl of the purified IgG samples were injected into UNcle sample holders (5 replicates for each sample). DLS measurement was performed at 25 °C (4 acquisition, 5 seconds each). Static light scattering (SLS) for $T_m$ and $T_{agg}$ measurement was carried out from 25 °C to 90 °C with a temperature increase of 0.3 °C per minute.

Please note that all the tested IgG samples (**1**-**10**) contained a ~5% defucosylated population. The biantennary samples (**2**-**4**) had ~10% bisecting glycoforms. The mono-antennary samples (**5**-**7**) had ~10% (F)M3 glycans due to the reversible activity of GnT-I. Sialylated samples (**4** & **10**) possessed about 1:1 mono- and di-sialylated populations. Please refer to **Figure 3** and **S12** for detailed glycans species and population distributions.

Binding assays between glycoengineered IgGs and Fc gamma receptor 1

This protocol is adapted from the AlphaLISA human FCGR binding kit manual provided by PerkinElmer. Briefly, serial dilutions of SPGR-engineered IgGs with 1X HiBlock buffer were prepared with the highest concentration at 1 mg/ml and the lowest concentration at 0.1 µg/ml. 10 µL of each diluted IgG samples were mixed with 10 µL 4X human FcγR1 solution and 20 µL Donor/acceptor beads solution into a white 96-well plate. The plate was then sealed and incubated at 25°C for 90 minutes without any light exposure. After the reaction, the fluorescence signal at 615 nm was determined using an EnVision Multimode Plate Reader (equipped with AlphaScreen module).

Computational modeling of IgG-protein A complex

Protein A homology model was constructed using the Swiss-Model server.(3) PDB structure 5H7B, which has 79.5% sequence identity with Protein A, was used as a template to construct the homology model.(4) A visual inspection of Protein A model illustrated 4 distinct IgG binding domains. PDB structure 5U4Y was used as a template to identify the spatial positioning of the full-length Protein A and the IgG.(5) The template structure contains only the B-domain of the protein A molecule. The spatial position of the B-domain helped us overlay the full-length protein-A molecule and allowed us to identify steric hindrances between other protein-A domains and the IgG molecule. Structure overlay and the movie illustrating clashes between the molecules was generated using Pymol (Molecular Graphics System, Version 2.0, Schrödinger LLC).

Plotting and graphic

Data plotting and curve fitting were done by using GraphPad Prism 8. Figures and cartoons were created by Adobe Illustrator.

### SI-Discussions

§1 Comparison between substrate immobilization and enzyme immobilization in SPGR

An alternative approach to conduct SPGR is immobilizing the glycoengineering enzymes on solid supports instead of the IgG substrates **(Figure S14)**.(6, 7) In this way, glycoengineering enzymes can be easily pulled out from the reaction pools for recovery. This could improve the efficiency of large-scale productions where the enzymes are re-charged in multiple batches of reactions. Furthermore, because substrates present the majority of the substance in the reactions, immobilizing the enzymes, instead of the substrate, could reduce the cost of immobilization in the production pipelines. However, there are also restrictions in this manner. For example, the selection of enzymes for the remodeling processes is much more limited in this case because the buffer of the IgG solution can not be swapped easily. In a remodeling process involving both glycosidases and glycosyltransferases—which prefer very different conditions (pH, cations)— the use of a generic working buffer would significantly compromise the enzyme activities. In addition, the preparation of resin-immobilized glycoengineering enzymes could be difficult. Whether immobilization affects their activity and substrate selectivity, as well as the immobilization protocol itself, remains to be investigated. Together, for the glycan remodeling that involves multiple glycoengineering enzymes, we believe the substrate-immobilization SPGR (this study) is preferred.

#### §2 Endoglycosidases for glycan remodeling

IgG glycan trimming can also be implemented through the GlcNAc residues in the chitobiose core. Endoglycosidases specifically cleave the β1-4 linkage between the GlcNAc residues in the chitobiose core.(8-11) They have been employed for chemoenzymatic glycan modification where the native glycans of targeted glycoprotein are first removed by endoglycosidases, followed by installing synthetic glycans back to the proteins using mutated endoglycosidases (glycosynthases).(12) Since re-building the chitobiose cores remains challenging, largely due to the lack of available mannosyl-transferases, we analyzed the activity of endoglycosidases on intact IgG but did not apply them in SPGR applications. Of the six tested enzymes, the candidate from *Streptococcus pyogenes* (known as Endo S) exhibited the highest conversion ratio on intact IgG ($CR_{50}$=0.7%, **Figure S15**). It has been known that endoglycosidases have different preferences for substrate structures. In agreement with reported studies, Endo S effectively liberates N-glycans from human IgG in SPGR.(9) On the other hand, the ones with higher specificity to high-mannose glycans, such as endoglycosidase D, did not show detectable IgG glycan conversion in our screening (**Table S1**).(13) An complete Endo S reaction led to the removal of glycan majority, giving a clean chromatogram as shown in **Figure 15**.

#### §3 *Candidatus omnitrophica* fucosidase has a wide spectrum on substrate selectivity

With the use of high enzyme concentration and long incubation time, we found that Fuc from *C. omnitrophica* functioned on all the IgG glycoforms. **(Figure S4c)** While we also observed decreased activity as the structural complexity of the glycans increased **(Figure S4d)**, the broad spectrum of substrate selectivity makes this enzyme an attractive tool for glycan remodeling on intact IgG. Opportunities to improve its activity through genetic engineering is worth to be investigated.

#### §4 Steric hindrance created by IgG immobilization reduces glycoengineering enzyme activities

IgG immobilization enables efficient washing and reaction-swapping processes in SPGR. However, we observed reduced enzyme activities on immobilized IgG compared to non-immobilized, free IgG **(Figure 16a)**. Such a "trade-off" partially results from the relatively limited surface area in heterogeneous reactions but could mainly be attributed to the increased steric hindrance created by IgG binding to protein A. It's reported that protein A binds to the IgG Fc at the interface between the CH2 and CH3 domains.(14) This binding region is not only closed to but also interacting with the Asn297 glycans.(15) Computational modeling of interactions between the full-length protein A (with four domains) and the Fc region indicates steric hindrance, specifically at the CH2 region **(Figure S16b-c)**. This could lead to reduced accessibility of the glycans by glycoengineering enzymes. This hypothesis of spatial hindrance and reduced accessibility is supported by the size effect of glycoengineering enzymes whose reduced activity correlates with their molecular weight. For example, the largest enzyme in our toolset, Gal from *S. pneumoniae* (231kD), showed an activity reduction of 74% when functions on immobilized IgG. A significant activity reduction was also found in Fuc despite its smaller size, an enzyme that enables chemistry at the very bottom of the glycan structure. In addition to steric hindrance, multiple IgG Fc regions could be interacting with the same protein A molecule, thus leading to a crowing effect and reduced enzymatic activity.

### SI-Tables

| | Enzyme | Source | Size (kD) | Targets (glycans with) | Conversion Ratio (%) 1 hour | Conversion Ratio (%) 24 hours | Vender & Cat # |
|---|---|---|---|---|---|---|---|
| ***Exoglycosidases*** | | | | | | | |
| V | Neuraminidase | *C. perfringens* | 43 | Terminal sialic acid | 66.4 ± 4.7 | 90.2 ± 1.3 | NEB P0720 |
| | Neuraminidase A | *A. ureafaciens* | 100 | Terminal sialic acid | 55 ± 3.5 | 56.5 ± 1.2 | NEB P0722 |
| | Neuraminidase S | *S. pneumoniae* | 74 | Terminal sialic acid | N.D. | 9.6 ± 1.6 | NEB P0743 |
| V | Galactosidase S | *S. pneumoniae* | 231 | Terminal galactose | 5.5 ± 2.2 | 51 ± 8.1 | NEB P0745 |
| | Galactosidase | *B. taurus* (testis) | 71 | Terminal galactose | 3.5 ± 0.9 | 3.7 ± 1.6 | NEB P0746 |
| | Galactosidase 1 | *H. sapiens* | 74 | Terminal galactose | N.D. | 3 ± 0.7 | R&D Systems 6464-GH |
| V | N-Acetylglucosaminidase S | *S. pneumoniae* | 125 | Terminal GlcNAc | 47 ± 1 | 82.9 ± 1.8 | NEB P0744 |
| | N-Acetylhexosaminidase F | *S. picatus* | 100 | Terminal GlcNAc | N.D. | N.D. | NEB P0721 |
| | Fucosidase | *B. Taurus* (kidney) | 52 | Terminal Fucose | N.D. | 1.2 ± 0.9 | NEB P0748 |
| V | Fucosidase O | *C. Omnitrophica* | 49 | Terminal Fucose | 1.3 ± 0.2 | 2.4 ± 1 | NEB P0749 |
| | Fucosidase | *Prunus dulcis* | 56 | Terminal Fucose | N.D. | N.D. | NEB P0769 |
| | Fucosidase | *C. meningosepticum* | 50 | Terminal Fucose | N.D. | N.D. | Sigma 344826 |
| | Fucosidase | *E. miricola* | 56 | Terminal Fucose | N.D. | N.D. | Sigma F6272 |
| | Fucosidase | *H. sapiens* | 51 | Terminal Fucose | N.D. | N.D. | R&D Systems 7039-GH |
| | Fucosidase | *T. maritima* | 54 | Terminal Fucose | N.D. | N.D. | R&D Systems 6556-GH |
| ***Endoglycosidases*** | | | | | | | |
| | Endoglycosidase A | *A. protophormia* | 69 | GlcNAc-GlcNAc linkage | N.D. | N.D. | Chemily G. EN01017 |
| | Endoglycosidase D | *S. pneumoniae* | 140 | GlcNAc-GlcNAc linkage | N.D. | N.D. | NEB P0742 |
| | Endoglycosidase F2 | *E. miricola* | 40 | GlcNAc-GlcNAc linkage | N.D. | N.D. | NEB P0772 |
| | Endoglycosidase F3 | *E. minicola* | 39 | GlcNAc-GlcNAc linkage | 17.1 ± 5.7 | 56.7 ± 13.4 | NEB P0771 |
| | Endoglycosidase M | *M. hiemalis* | 85 | GlcNAc-GlcNAc linkage | N.D. | N.D. | TCI A1651 |
| V | Endoglycosidase S | *S. pyogenes* | 136 | GlcNAc-GlcNAc linkage | 39.9 ± 1.9 | 68.4 ± 5.2 | NEB P0741 |
| ***Glycosyltransferases*** | | | | | | | |
| | α2-6 Sialyltransferase | *P. damsela* | 59 | Terminal Galactose | N.D. | N.D. | Sigma S2076 |
| | α2-6 Sialyltransferase | *P. multocida* | 46 | Terminal Galactose | N.D. | N.D. | Sigma S1951 |
| V | α2-6 Sialyltransferase (ST6Gal1) | *H. sapiens* | 44 | Terminal Galactose | 7.8 ± 1.6 | 29.3 ± 0.4 | Sigma SAE0090 |
| | α2-6 Sialyltransferase (ST6Gal2) | *H. sapiens* | 33 | Terminal Galactose | N.D. | 1.6 ± 1.2 | R&D Systems 8330-GT |
| | α2-6 Sialyltransferase (ST6GalNAc4) | *H. sapiens* | 31 | Terminal Galactose | N.D. | 1.2 ± 0.4 | R&D Systems 6876-GT |
| | α2-3 Sialyltransferase (ST3-Gal1) | *H. sapiens* | 33 | Terminal Galactose | N.D. | 2.1 ± 0.4 | R&D Systems 6905-GT |
| | α2-3 Sialyltransferase (ST3-Gal2) | *H. sapiens* | 35 | Terminal Galactose | N.D. | 1.2 ± 1.1 | R&D Systems 7275-GT |
| V | β1-4 Galactosyltransferase 1 | Homo sapiens | 40 | Terminal GlcNAc | 7 ± 1.8 | 22.2 ± 1.1 | Sigma SAE0093 |
| | β1-4 Galactosyltransferase | *B. Taurus* (milk) | 45 | Terminal GlcNAc | 3.1 ± 1 | 23 ± 1.2 | Sigma G5507 |
| V | N-Acetylglucosaminyltransferase 1 | *H. sapiens* | 48 | (F)M3 glycoforms | 8.4 ± 2.6* | 17.6 ± 0.2* | R&D Systems 8334-GT |
| V | N-Acetylglucosaminyltransferase 3 | *H. sapiens* | 58 | (F)A2 glycoforms | N.D. | 6 ± 2.1 | R&D Systems 7359-GT |
| V | N-Acetylglucosaminyltransferase 5 | *H. sapiens* | 65 | (F)A2/(F)A3 glycoforms | N.D. | N.D. | R&D Systems 5469-GT |
| V | Fucosyltransferase 8 | *H. sapiens* | 64 | (F)M3 glycoforms | 1.7± 0.6 | 9.2 ± 2.1 | R&D Systems 5768-GT |

**Table S1**. Activity screening of glycoengineering enzymes. Enzyme activity was determined by using SPGR protocol with 1mg human serum IgG (66.7 µM) and glycoengineering enzyme (0.25 µM) for 1 or 24 hours. HILIC-MS was employed for characterizing glycan structures and quantification. N.D. = not detectable. V= enzymes showed the highest activity in the screening (selected for SPGR). 0.1 ml Protein A resin was used for immobilization and the final reaction volume was 0.1 ml for all the reactions. Reactions in this screening were conducted using the working conditions suggested by the vendors without further optimization. *Glycoengineered IgG was used as the substrate for the screening.

| | Exp. | Enzyme (Amount) | Substrate | Buffer | Mol % (E-to-S) | Weight % (E-to-S) | Reaction Time | Temp. | pH | Cation |
|---|---|---|---|---|---|---|---|---|---|---|
| | Fig S1A | Neuraminidase (2 µg) | Human serum IgG | 50 mM Sodium acetate | 0.7% | 0.2% | 1h | **Variable** | 5.5 | $Ca^{2+}$ |
| | Fig S1B | Neuraminidase | Human serum IgG | 50 mM Sodium acetate | **Variable** | **Variable** | 1h | 42°C | 5.5 | $Ca^{2+}$ |
| | Fig S1C | Neuraminidase (16 µg) | Human serum IgG | 50 mM Sodium acetate | 5.6% | 1.6% | **Variable** | 42°C | 5.5 | $Ca^{2+}$ |
| | Fig 2 | Neuraminidase (32 µg) | Human serum IgG | 50 mM Sodium acetate | 11.2% | 3.2% | 4h | 42°C | 5.5 | $Ca^{2+}$ |
| | Fig S2A | Galactosidase S (12.5 µg) | Human serum IgG | 50 mM Sodium acetate | 0.75% | 1.25% | 1h | **Variable** | 5.5 | $Ca^{2+}$ |
| | Fig S2B | Galactosidase S | Human serum IgG | 50 mM Sodium acetate | **Variable** | **Variable** | 1h | 37°C | 5.5 | $Ca^{2+}$ |
| | Fig S2C | Galactosidase S (75 µg) | Human serum IgG | 50 mM Sodium acetate | 4.5% | 7.5% | **Variable** | 37°C | 5.5 | $Ca^{2+}$ |
| | Fig 2 | Galactosidase S (37.5 µg) | Human serum IgG | 50 mM Sodium acetate | 2.25% | 3.75% | 16h | 37°C | 5.5 | $Ca^{2+}$ |
| | Fig S3A | N-Acetylglucosaminidase S (3 µg) | Human serum IgG | 50 mM Sodium acetate | 0.37% | 0.3% | 1h | **Variable** | 5.5 | $Ca^{2+}$ |
| | Fig S3B | N-Acetylglucosaminidase S | Human serum IgG | 50 mM Sodium acetate | **Variable** | **Variable** | 1h | 42°C | 5.5 | $Ca^{2+}$ |
| | Fig S3C | N-Acetylglucosaminidase S (25 µg) | Human serum IgG | 50 mM Sodium acetate | 3.2% | 2.5% | **Variable** | 42°C | 5.5 | $Ca^{2+}$ |
| | Fig 2 | N-Acetylglucosaminidase S (35 µg) | Human serum IgG | 50 mM Sodium acetate | 4.2% | 3.5% | 4h | 42°C | 5.5 | $Ca^{2+}$ |
| | Fig S4A | Fucosidase O (1.32 mg) | FM3 IgG | 50 mM Sodium acetate | 400% | 132% | 3-days | 37°C | 4.5 | None |
| | Fig S4C | Fucosidase O (0.33 mg) | Human serum IgG | 50 mM Sodium acetate | 100% | 33% | 5-days | 37°C | 4.5 | None |
| | Fig S15A | Endoglycosidase S (27 µg) | Human serum IgG | 50 mM Sodium acetate | 3% | 2.7% | 4h | 42°C | 5.5 | $Ca^{2+}$ |
| | Fig S15B | Endoglycosidase S (3.4 µg) | Human serum IgG | 50 mM Sodium acetate | 0.37% | 0.34% | 1h | **Variable** | 5.5 | $Ca^{2+}$ |
| | Fig S15C | Endoglycosidase S | Human serum IgG | 50 mM Sodium acetate | **Variable** | **Variable** | 1h | 42°C | 5.5 | $Ca^{2+}$ |
| | Fig S15D | Endoglycosidase S (13.6 µg) | Human serum IgG | 50 mM Sodium acetate | 1.5% | 1.36% | **Variable** | 42°C | 5.5 | $Ca^{2+}$ |
| | Fig S5A | α2-6 Sialyltransferase (1.1 µg) | Human serum IgG | 25 mM Tris-HCl | 0.37% | 0.11% | 24h | **Variable** | 7.5 | $Ca^{2+}$, $Mn^{2+}$ |
| | Fig S5B | α2-6 Sialyltransferase (1.1 µg) | Human serum IgG | 25 mM Tris-HCl | 0.37% | 0.11% | 24h | 37°C | **Variable** | $Ca^{2+}$, $Mn^{2+}$ |
| | Fig S5C | α2-6 Sialyltransferase (1.1 µg) | Human serum IgG | 25 mM Tris-HCl | 0.37% | 0.11% | 24h | 37°C | 7.5 | **Variable** |
| | Fig S5D | α2-6 Sialyltransferase | Human serum IgG | 25 mM Tris-HCl | **Variable** | **Variable** | 1h | 37°C | 7.5 | $Mg^{2+}$ |
| | Fig S5E | α2-6 Sialyltransferase (44 µg) | Human serum IgG | 25 mM Tris-HCl | 14.8% | 4.4% | **Variable** | 37°C | 7.5 | $Mg^{2+}$ |
| | Fig 2 | α2-6 Sialyltransferase (15 µg) | Human serum IgG | 25 mM Tris-HCl | 5% | 1.5% | 16h | 37°C | 7.5 | $Mg^{2+}$ |
| | Fig S6A | β1-4 Galactosyltransferase 1 (10 µg) | Human serum IgG | 25 mM Tris-HCl | 3.7% | 1% | 1h | **Variable** | 7 | $Na^{+}$, $Mn^{2+}$ |
| | Fig S6B | β1-4 Galactosyltransferase 1 (8 µg) | Human serum IgG | 25 mM Tris-HCl | 3% | 0.8% | 2h | 50°C | **Variable** | $Na^{2+}$, $Mn^{2+}$ |
| | Fig S6C | β1-4 Galactosyltransferase 1 (4 µg) | Human serum IgG | 25 mM Tris-HCl | 1.5% | 0.4% | 3h | 50°C | 7 | **Variable** |
| | Fig S6D | β1-4 Galactosyltransferase 1 | Human serum IgG | 25 mM Tris-HCl | **Variable** | **Variable** | 1h | 50°C | 7 | $Mn^{2+}$ |
| | Fig S6E | β1-4 Galactosyltransferase 1 (30 µg) | Human serum IgG | 25 mM Tris-HCl | 11.2% | 3% | **Variable** | 50°C | 7 | $Mn^{2+}$ |
| | Fig 2 | β1-4 Galactosyltransferase 1 (25 µg) | Human serum IgG | 25 mM Tris-HCl | 9.4% | 2.5 | 16 | 50°C | 7 | $Mn^{2+}$ |
| | Fig S8A | N-Acetylglucosaminyltransferase 1 (1.2 µg) | FM3 IgG | 20 mM HEPES | 0.37% | 0.12% | 4h | **Variable** | 7 | $Na^{+}$, $Mn^{2+}$ |
| | Fig S8B | N-Acetylglucosaminyltransferase 1 (1.2 µg) | FM3 IgG | 20 mM HEPES | 0.37% | 0.12% | 4h | 37°C | **Variable** | $Na^{+}$, $Mn^{2+}$ |
| | Fig S8C | N-Acetylglucosaminyltransferase 1 (2.4 µg) | FM3 IgG | 20 mM HEPES | 0.74% | 0.24% | 2h | 30°C | 7.5 | **Variable** |
| | Fig S8D | N-Acetylglucosaminyltransferase 1 | FM3 IgG | 20 mM HEPES | **Variable** | **Variable** | 1h | 30°C | 7.5 | $Na^{+}$, $Ca^{2+}$, $Mg^{2+}$, $Mn^{2+}$ |
| | Fig S8E | N-Acetylglucosaminyltransferase 1 (15 µg) | FM3 IgG | 20 mM HEPES | 4.5% | 1.5% | **Variable** | 30°C | 7.5 | $Na^{+}$, $Ca^{2+}$, $Mg^{2+}$, $Mn^{2+}$ |
| | Fig S8F | N-Acetylglucosaminyltransferase 1 (30 µg) | FM3 IgG | 20 mM HEPES | **Variable** | **Variable** | **Variable** | 30°C | 7.5 | $Na^{+}$, $Ca^{2+}$, $Mg^{2+}$,$Mn^{2+}$ |
| | Fig S7 | N-Acetylglucosaminyltransferase 1 (20 µg) | FM3 IgG | 20 mM HEPES | 6% | 2% | 4 | 30°C | 7.5 | $Na^{+}$, $Ca^{2+}$, $Mg^{2+}$, $Mn^{2+}$ |
| | Fig S9A | N-Acetylglucosaminyltransferase 3 (9 µg) | Human serum IgG | 20 mM HEPES | 1.5% | 0.9% | 24h | **Variable** | 7 | $Na^{+}$, $Mn^{2+}$ |
| | Fig S9B | N-Acetylglucosaminyltransferase 3 (9 µg) | Human serum IgG | 20 mM HEPES or MES | 1.5% | 0.9% | 24h | 37°C | **Variable** | $Na^{+}$, $Mn^{2+}$ |
| | Fig S9C | N-Acetylglucosaminyltransferase 3 (9 µg) | Human serum IgG | 20 mM MES | 1.5% | 0.9% | 24h | 30°C | 6.5 | **Variable** |
| | Fig S9D | N-Acetylglucosaminyltransferase 3 | Human serum IgG | 20 mM MES | **Variable** | **Variable** | 1h | 30°C | 6.5 | $Na^{+}$, $Mn^{2+}$ |
| | Fig S9E | N-Acetylglucosaminyltransferase 3 (45 µg) | Human serum IgG | 20 mM MES | 7.5% | 4.5% | **Variable** | 30°C | 6.5 | $Na^{+}$, $Mn^{2+}$ |
| | Fig 2 | N-Acetylglucosaminyltransferase 3 (30 µg) | Human serum IgG | 20 mM MES | 5% | 3% | 16h | 30°C | 6.5 | $Na^{+}$, $Mn^{2+}$ |
| | Fig S10 | N-Acetylglucosaminyltransferase 5 (33 µg) | Human serum IgG | 20 mM HEPES | 7.5% | 3.3% | 24h | 30°C | 7.5 | $Na^{+}$, $Mn^{2+}$ |
| | Fig S11 | Fucosyltransferase (60 µg) | Defucosylated IgG | 100 mM MES | 14% | 6% | 5 days | 37°C | 7 | $Na^{+}$ |

**Table S2.** Reaction conditions used for SPGR reactions in this work. Protein A resin (0.1 ml) was used for IgG (1 mg) immobilization for all the reactions. The final reaction volume was 0.1 ml. (F)M3 IgG was prepared from human serum IgG by applying sequential glycan remodeling using SPGR as described in this study. Cation ion concentration: 5 mM for $Ca^{2+}$ ($CaCl_2$), 50 mM for $Na^{+}$ (NaCl) and 10 mM for $Mg^{2+}$ and $Mn^{2+}$ ($MgCl_2$, $MnCl_2$).

| | Serum IgG | Endo S-treated | Fuc-treated | Featured terminal (Bi-antennary) | | | | Mono-antennary | | | Bisecting | | |
|---|---|---|---|---|---|---|---|---|---|---|---|---|---|
| | | | | 1 | 2 | 3 | 4 | 5 | 6 | 7 | 8 | 9 | 10 |
| **Hydrodynamic diameter (nm)** | 13.3±1.2 | -- | -- | 11.5±0.5 | 17.7±2 | 13.6±0.4 | 11.8±0.6 | 14.8±0.7 | 14.2±0.9 | 13.3±0.9 | 17.4±1.2 | 12.8±0.4 | 18±2.4 |
| **Melting temperature (°C)** | 66.7±0.7 | 56.4±1.4 | -- | 63.4±0.4 | 64.8±0.7 | 66.3±0.3 | 66.1±0.2 | 61.4±0.5 | 62.9±0.7 | 63±0.4 | 64.8±1.1 | 64.5±0.3 | 65±0.7 |
| **Aggregation temperature (°C)** | 62±1.7 | 48.6±1.3 | -- | 59±2.3 | 63.3±1 | 66.1±0.5 | 64.3±1.1 | 62.2±0.5 | 62.2±1.2 | 63±0.9 | 60.9±1.2 | 63.3±0.4 | 64.9±0.6 |
| **FcγR I binding ($EC_{50}$, $10^{-7}$ g/ml)** | 1.9±0.4 | -- | 1.6±0.4 | 2.2±0.7 | 2.3±0.5 | 1.7±0.5 | 2±0.2 | 2.3±0.3 | 2.5±0.2 | 2.4±0.3 | 0.8±0.1 | 0.9±0.1 | 1.2±0.3 |

**Table S3. Physical and biochemical properties of SPGR-engineered IgGs**

| | Enzyme | Donor | Substrate glycan species (Acceptor) (excluding low-abundant, non-detectable glycoforms) |
|---|---|---|---|
| | Neuraminidase | N.A. | (F)A2(B)G2S2, (F)A2(B)G2S1, (F)A2(B)G1S1 |
| | Galactosidase S | N.A. | (F)A2(B)G2S1, (F)A2(B)G2, (F)A2(B)G1 |
| | N-Acetylglucosaminidase S | N.A. | (F)A2(B)G1, (F)A2(B) |
| | Fucosidase O | N.A. | All fucosylated IgG glycans |
| | Endoglycosidase S | N.A. | All IgG glycans |
| | α2-6 Sialyltransferase | Cytidine-5’-monophospho-N-acetylneuraminic acid (CMP-NANA) | (F)A2(B)G2, (F)A2(B)G1, (F)A2(B)G2S1 |
| | β1-4 Galactosyltransferase 1 | Uridine 5’-diphosphogalactose (UDP-Gal) | (F)A2(B), (F)A2(B)G1 |
| | N-Acetylglucosaminyltransferase 1 | Uridine 5’-diphospho-N-acetylglucosamine (UDP-GlcNAc) | (F)M3 (glycoengineered IgG) |
| | N-Acetylglucosaminyltransferase 3 | Uridine 5’-diphospho-N-acetylglucosamine (UDP-GlcNAc) | (F)A2, (F)A2G1 |
| | Fucosyltransferase | Guanosine 5’-diphospho-β-L-fucose (GDP-Fuc) | M3 (glycoengineered IgG) |

**Table S4.** Saccharide donor and acceptors in each enzymatic reaction. The substrate glycan species (acceptors) were listed based on our observation in this study.

**SI-Figures**

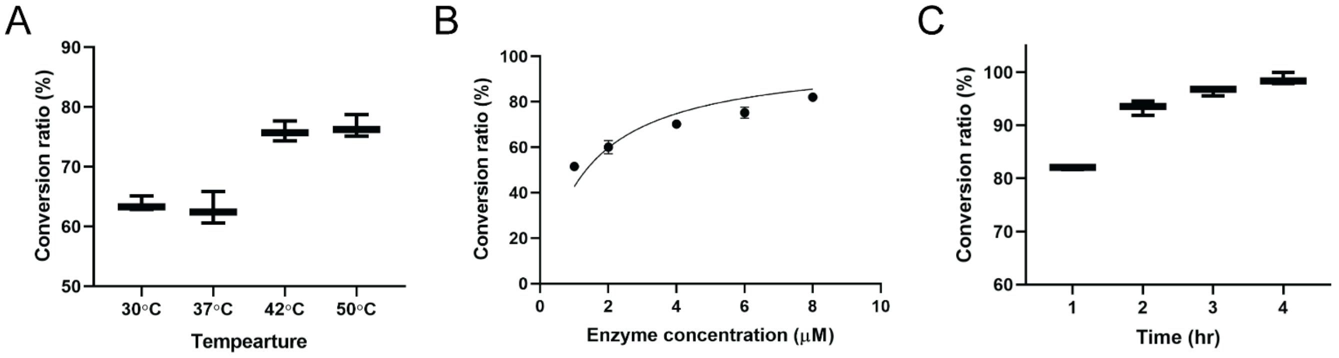

**Figure S1.** Characterization of Neuraminidase (*C. perfringens*) activity and its working condition optimization. (A) Temperature optimization. (B) Dose-dependent experiment. (C) Time-course study. Error bars: mean, maximum and minimum values. Refer to **Table S2** for reaction conditions.

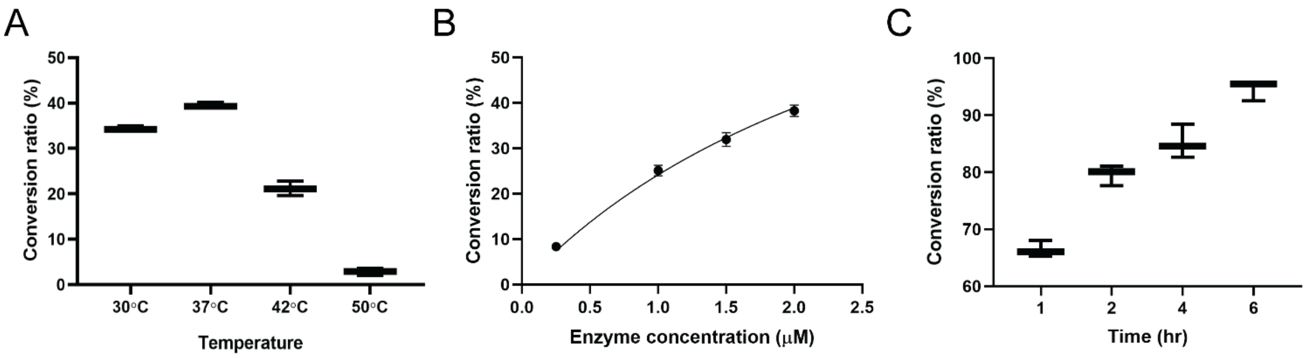

**Figure S2.** Characterization of Galactosidase S (*S. pneumoniae*) activity and its working condition optimization. (A) Temperature optimization. (B) Dose-dependent experiment. (C) Time-course study. Error bars: mean, maximum and minimum values. Refer to **Table S2** for reaction conditions.

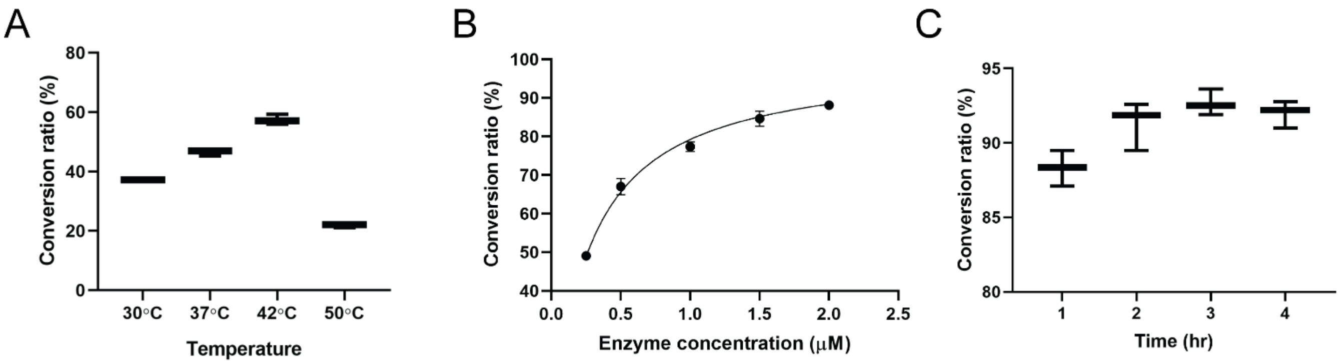

**Figure S3.** Characterization of *N*-Acetylglucosaminidase S (*S. pneumoniae*) activity and its working condition optimization. (A) Temperature optimization. (B) Dose-dependent experiment. (C) Time-course study. Error bars: mean, maximum and minimum values. Refer to **Table S2** for reaction conditions.

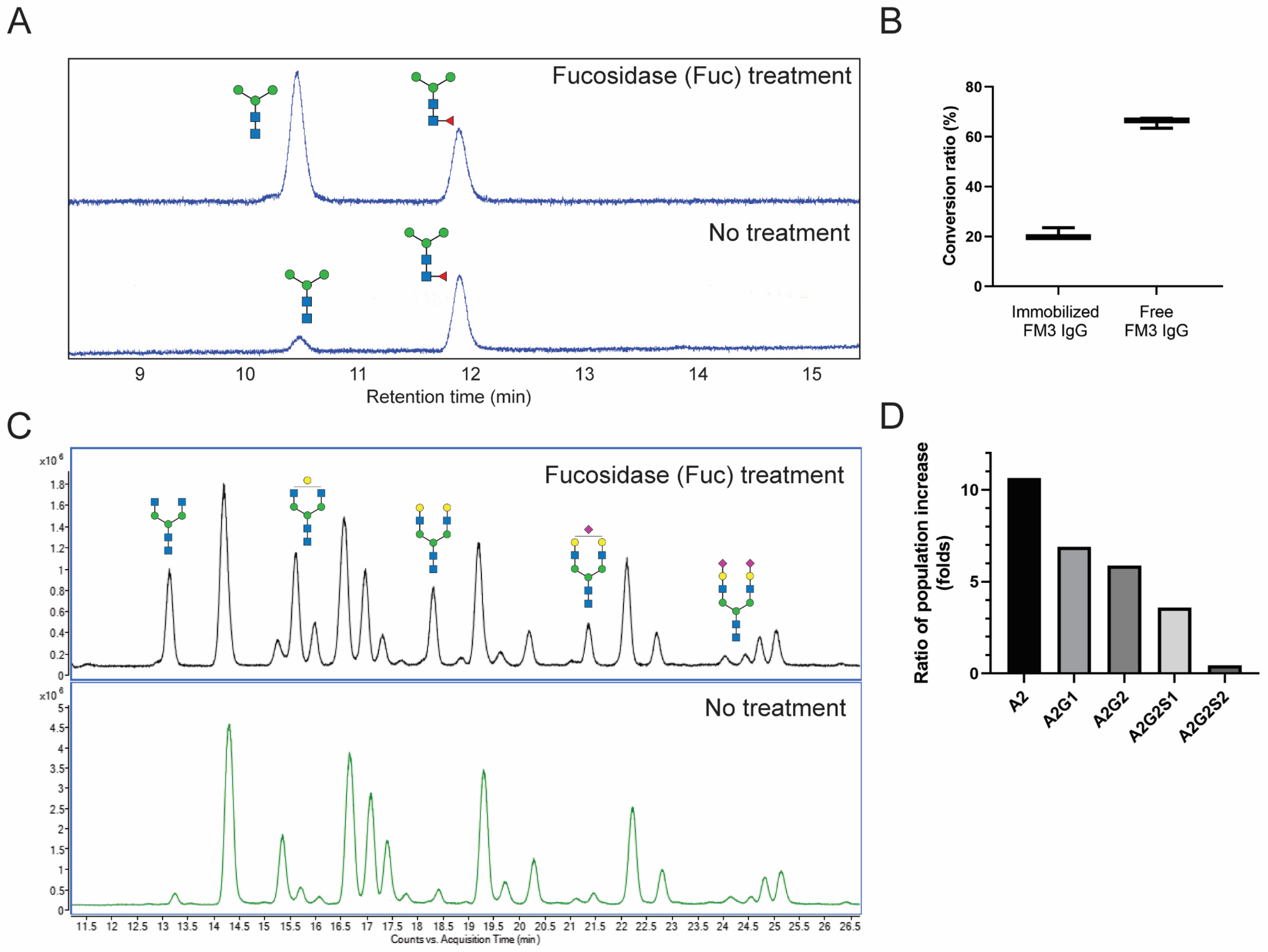

**Figure S4.** Characterization of fucosidase (*Candidatus Omnitrophica*) activity on intact IgG. (A) A 3-days reaction with intact IgG bearing (F)M3 glycans revealed the activity of fucosidase. The (F)M3 glycoforms were prepared using SPGR. (B) Comparison of conversion ratio between reactions with immobilized IgG and free IgG. (C) A 5-days reaction with intact IgG (no immobilization) revealed the broad substrate spectrum of the enzyme. Defucosylated populations were increased (highlighted with glycan images). See supplementary discussion for more details. (D) Fucosidase activity decreased as the structural complexity of glycans increased. Refer to **Table S2** for reaction conditions.

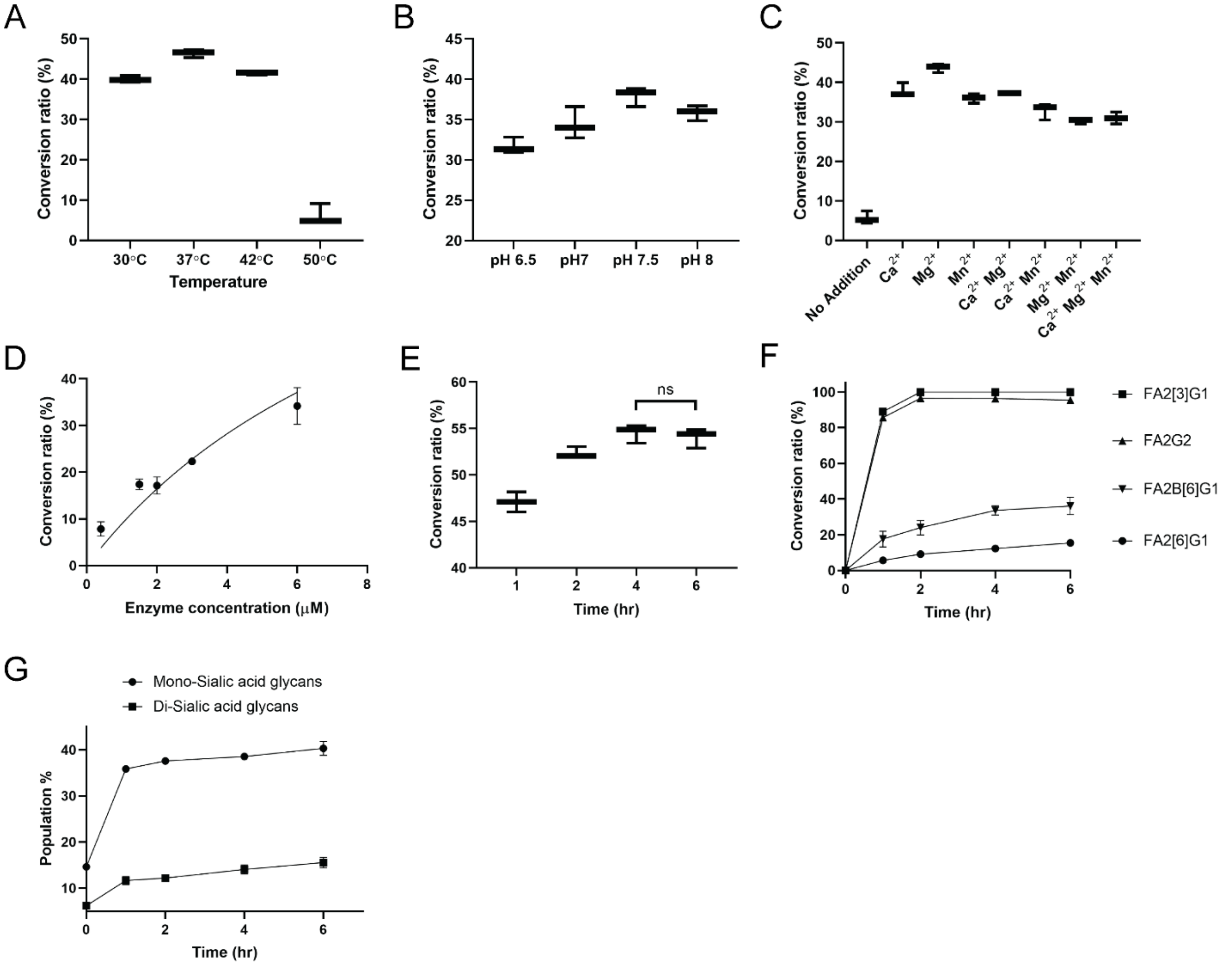

**Figure S5.** Characterization of α2-6 Sialyltransferase (*H. sapiens*) activity and its working condition optimization. (A) Temperature optimization. (B) pH optimization. (C) Cation optimization. (D) Dose-dependent experiment. (E) Time-course study. Conversion ratio calculations for α2-6 Sialyltransferase reactions were based on the consumption of non-sialylated glycans with terminal galactoses. (F) Comparison of enzyme activity between different IgG glycoforms. The data was collected from the time-course experiments. (G) Comparison of enzyme activity between the installation of 1st and 2nd sialic acid. Refer to **Table S2** for reaction conditions.

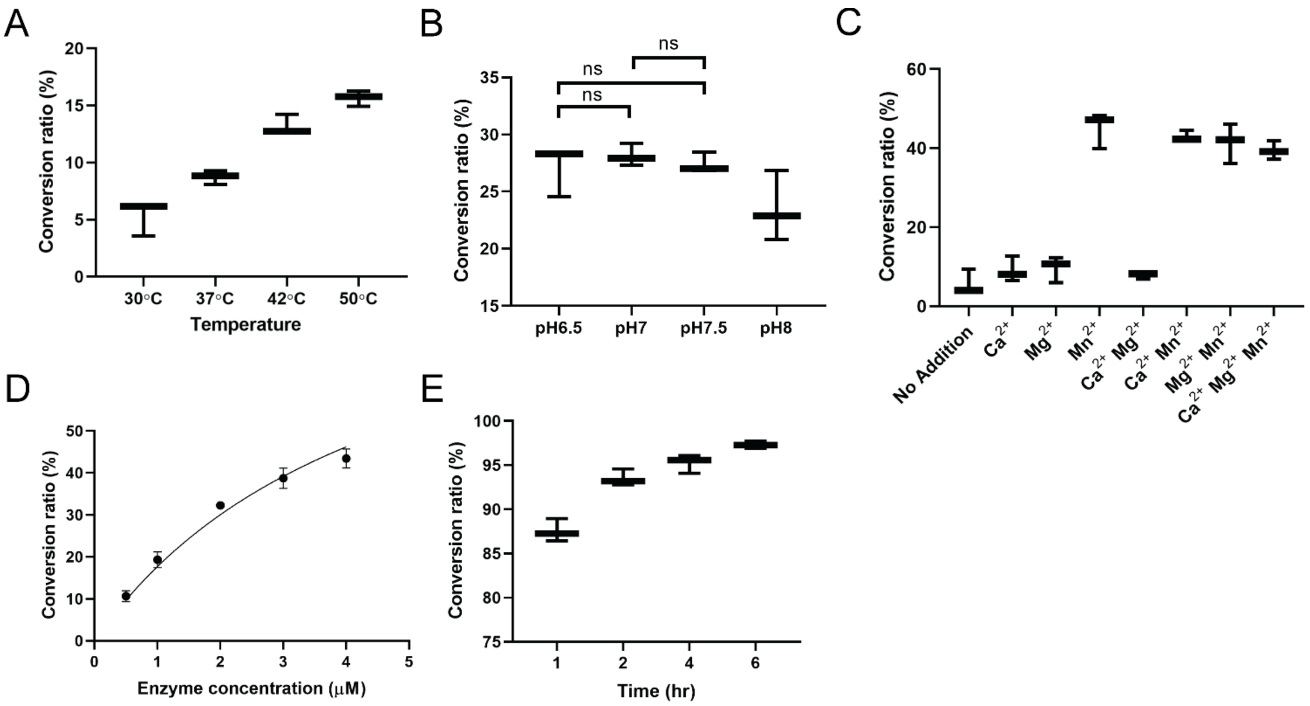

**Figure S6.** Characterization of β1-4 Galactosyltransferase 1 (*H. sapiens*) activity and its working condition optimization. (A) Temperature optimization. (B) pH optimization. (C) Cation optimization. (D) Dose-dependent experiment. (E) Time-course study. Error bars: mean, maximum and minimum values. Refer to **Table S2** for reaction conditions.

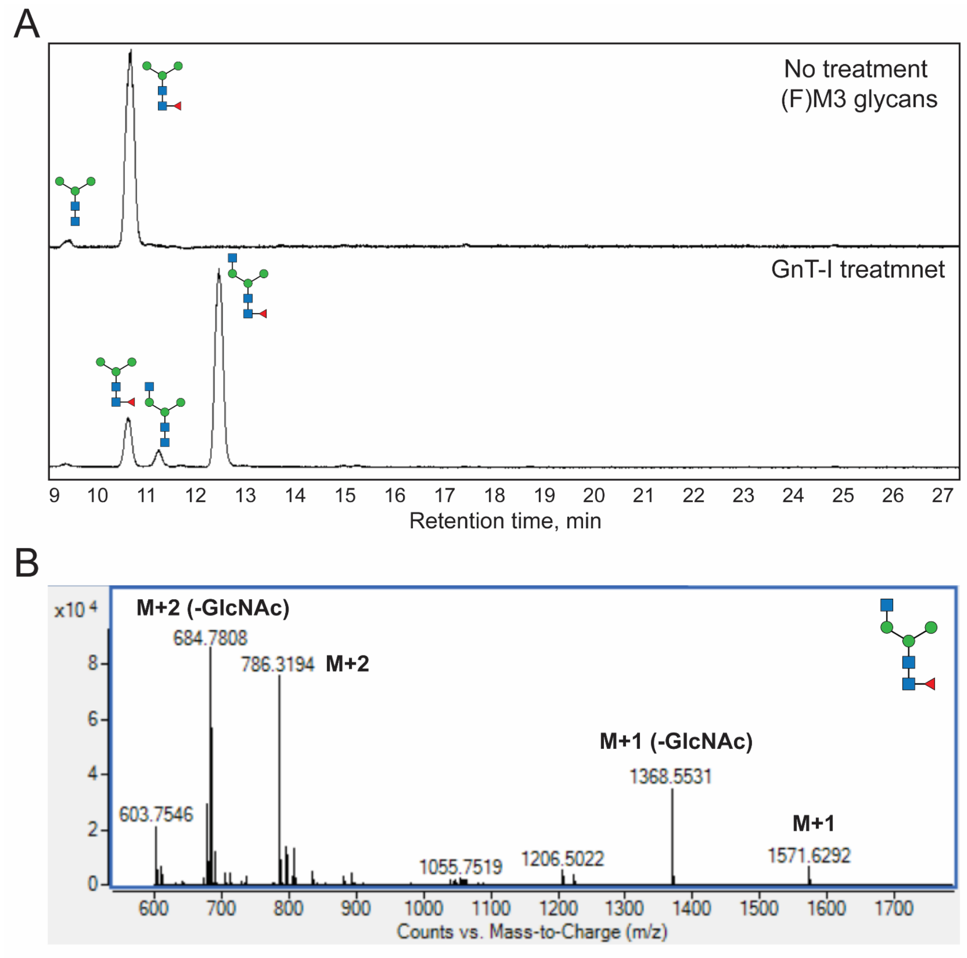

**Figure S7.** Characterization of *N*-Acetylglucosaminyltransferases I (GnT-I, MGAT1) activity. (A) Chromatographic analyses revealed GnT-I activity on (F)M3 glycans prepared by SPGR. (B) Mass spectrometry confirmed the formation of (F)A1[3] glycans. Glycan samples for the mass spectrometry analyses were labeled by RapiFlour-MS probe (MW=312.4) from Waters. On-column cleavage of terminal GlcNAc residues was often observed in HPLC-MS analyses. The IgG substrate bearing (F)M3 glycans was prepared by SPGR.

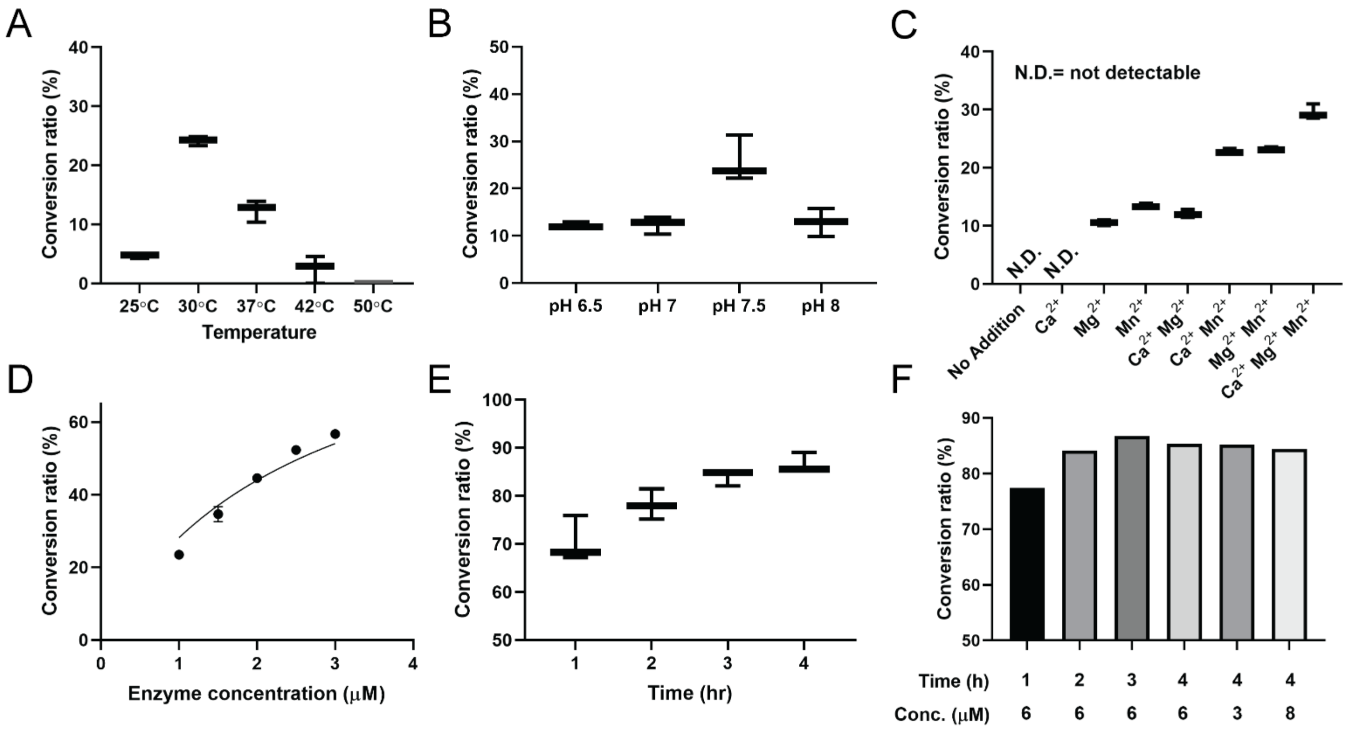

**Figure S8.** Characterization of *N*-Acetylglucosaminyltransferase 1 (GnT-I, *H. sapiens*) activity and its working condition optimization. (A) Temperature optimization. (B) pH optimization. (C) Cation optimization. (D) Dose-dependent experiment. (E) Time-course study. (F) The conversion ratio reached a plateau at ~ 85%. Error bars: mean, maximum and minimum values. Refer to **Table S2** for reaction conditions.

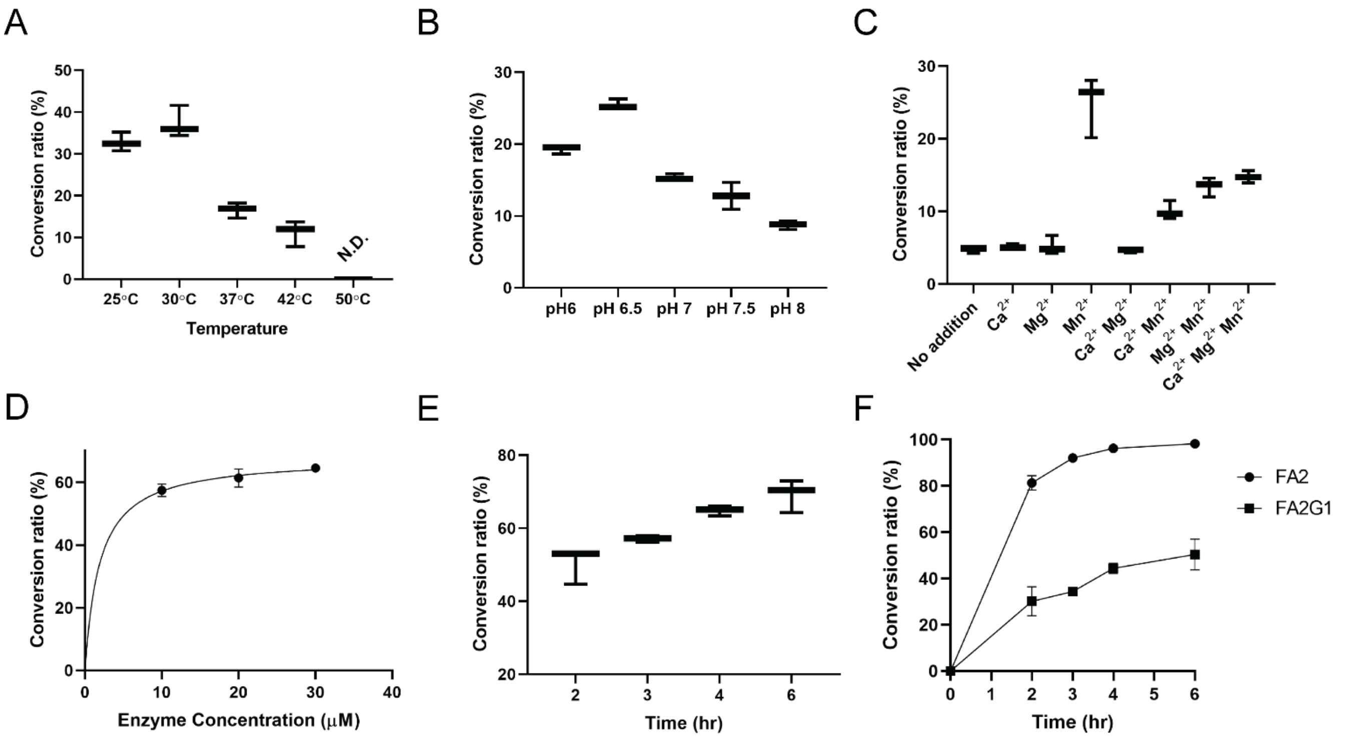

**Figure S9.** Characterization of *N*-Acetylglucosaminyltransferase 3 (GnT-III, *H. sapiens*) activity and its working condition optimization. (A) Temperature optimization. (B) pH optimization. (C) Cation optimization. (D) Dose-dependent experiment. (E) Time-course study. (F) Comparison of enzyme activity (in conversion ratio) between different IgG glycoforms. The data was collected from the time-course experiments. Refer to **Table S2** for reaction conditions.

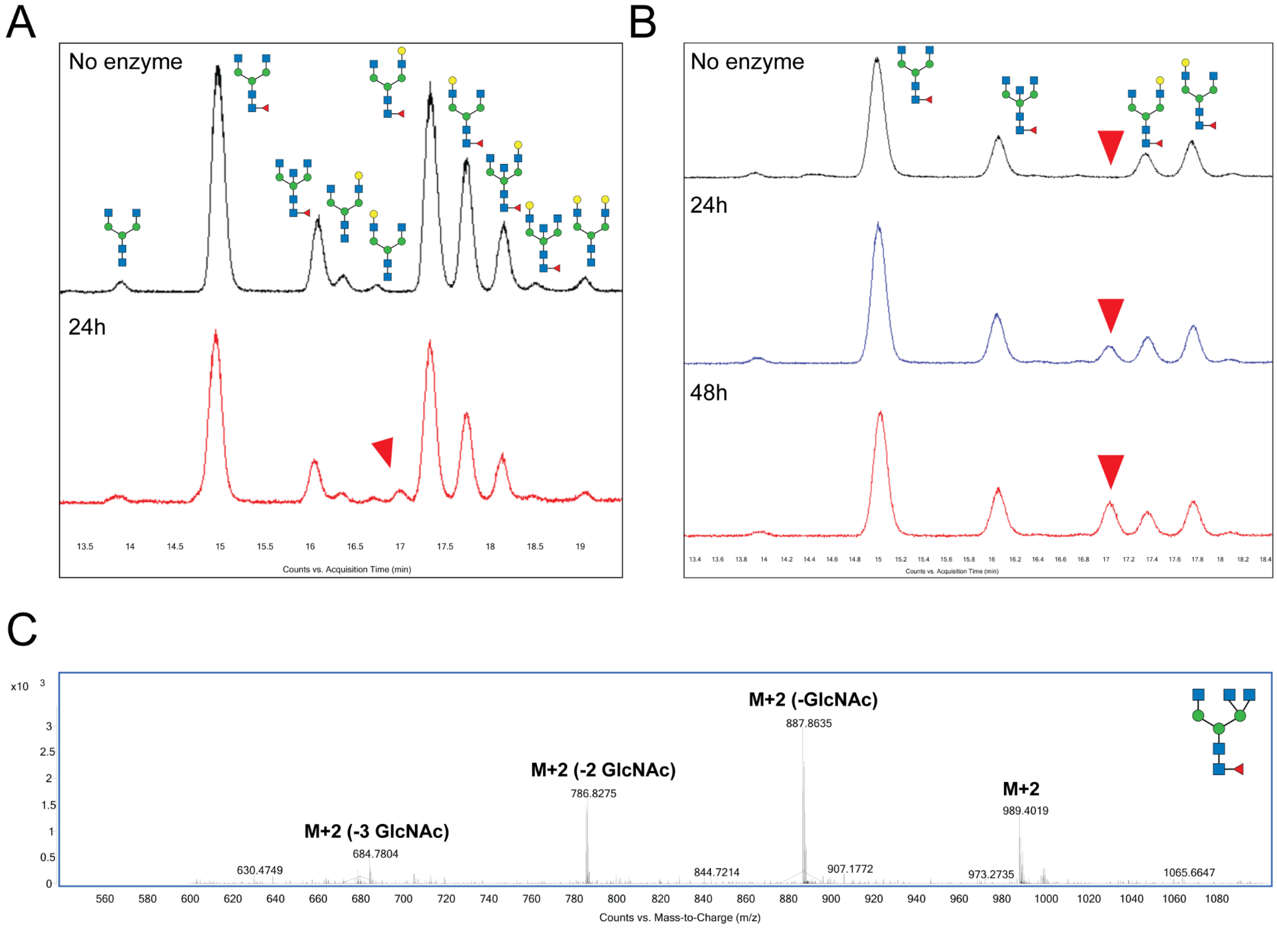

**Figure S10.** *In vitro* activity of *N*-Acetylglucosaminyltransferase 5 (GnT-V, *H. sapiens*) on intact human serum IgG immobilized on protein A resin. (A) GnT-V reaction with native human IgG. A new peak (indicated by red arrowhead) was found after a 24 hours reaction. (B) GnT-V reaction with glycoengineered human IgG. Human serum IgG was first treated with galactosidase in order to minimize the overlapping between the product signal and the FA2G1 glycan signals. A signal increase in the newly found peak was observed when the incubation time was increased. (C) Mass spectrometry analysis of the newly formed peak after the GnT-V reaction. The molecular weight (reported in m/z) of FA3 glycan was confirmed. The fragments resulting from on-column cleavage of GlcNA also supported that this analyte contains three terminal GlcNAc. The newly formed FA3 glycan has a 1-minute shift in retention time from that of FA2B glycan, despite they share the same molecular weight. Glycan samples for the mass spectrometry analyses showing here were labeled by RapiFlour-MS probe from Waters (MW=312.4).

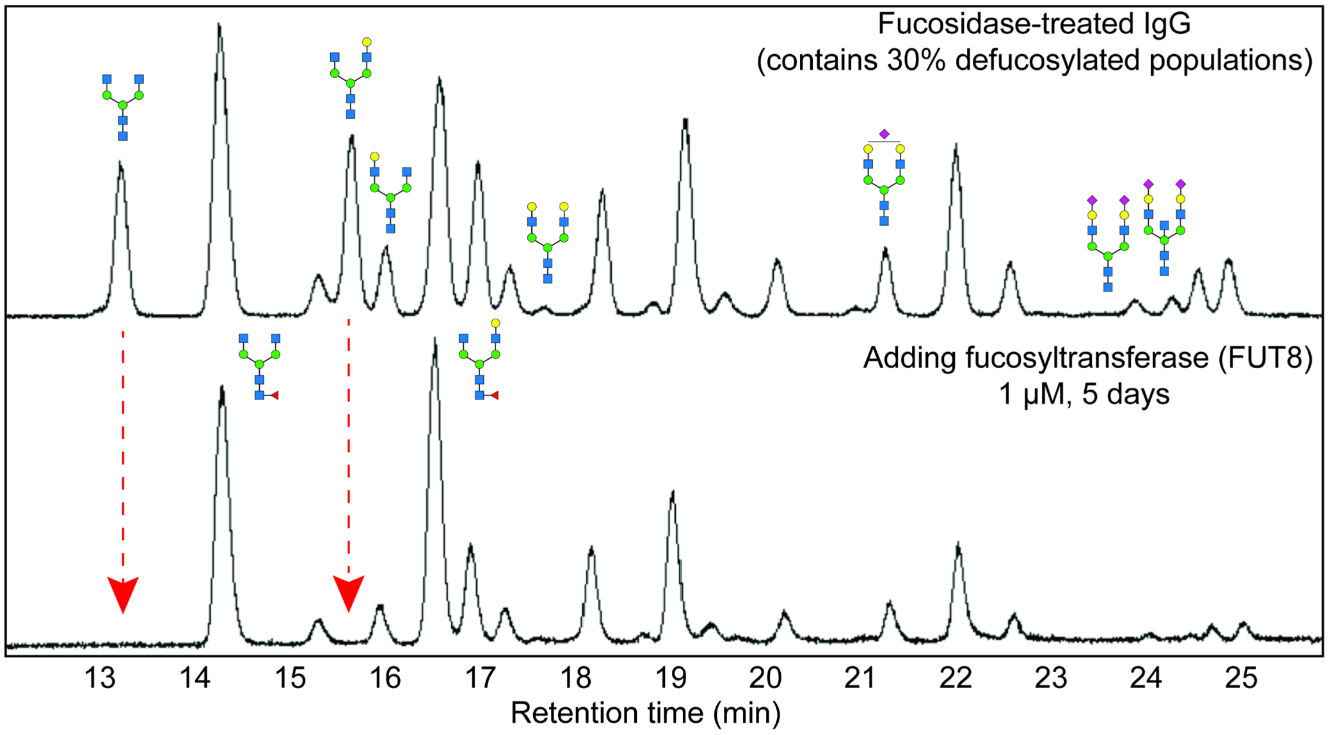

**Figure S11.** Characterization of fucosyltransferase (*Homo sapiens,* FUT8) activity. Serum IgG was treated with fucosidase (100 mol%) for 5 days. The resulting product (30% of the IgG population was defucosylated) was then used for the FUT8 reaction. Red arrows indicate the glycoforms consumed by FUT8. Refer to **Table S2** for reaction conditions.

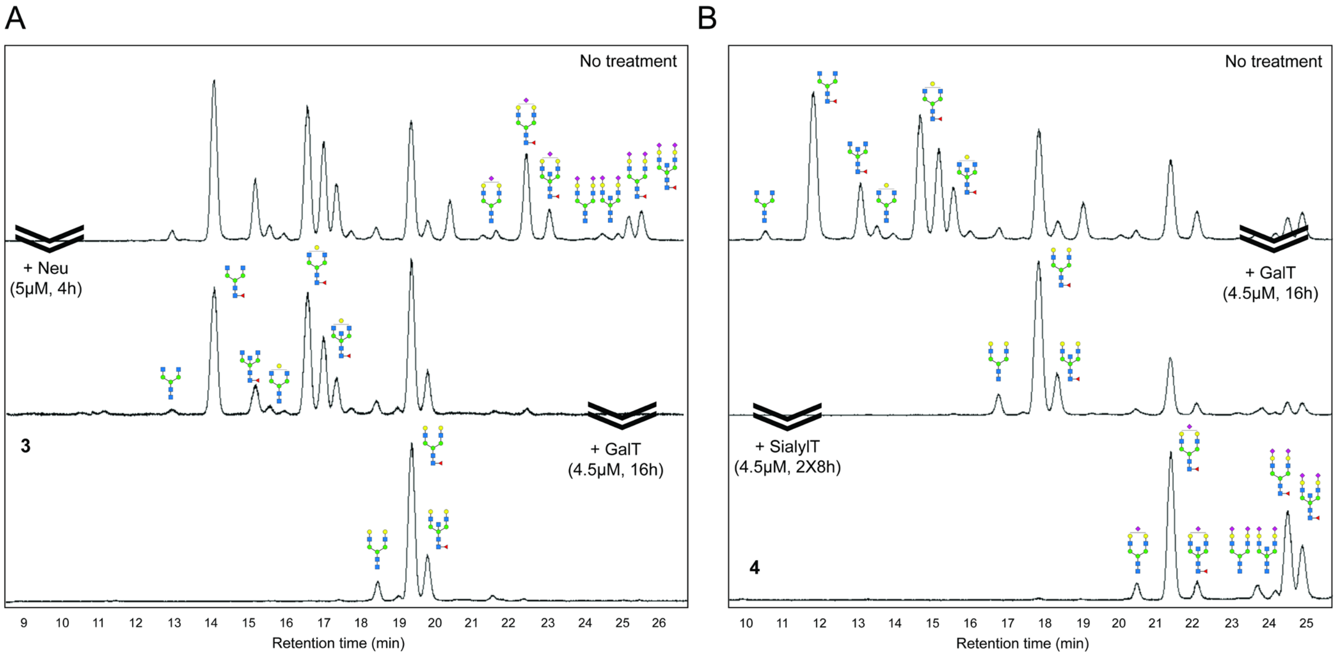

**Figure S12.** Chromatogram of fluorescently labeled glycans collected from IgG undergoing successive glycan remodeling using SPGR. (A) Harmonization of terminal residues into galactose. (B) Harmonization of terminal residues into sialic acid. The reactions were conducted on 1 mg human serum IgG immobilized on 0.1 ml protein A resin. Please refer **Figure 4** for the sample numbering and **Figure 1e** for working conditions.

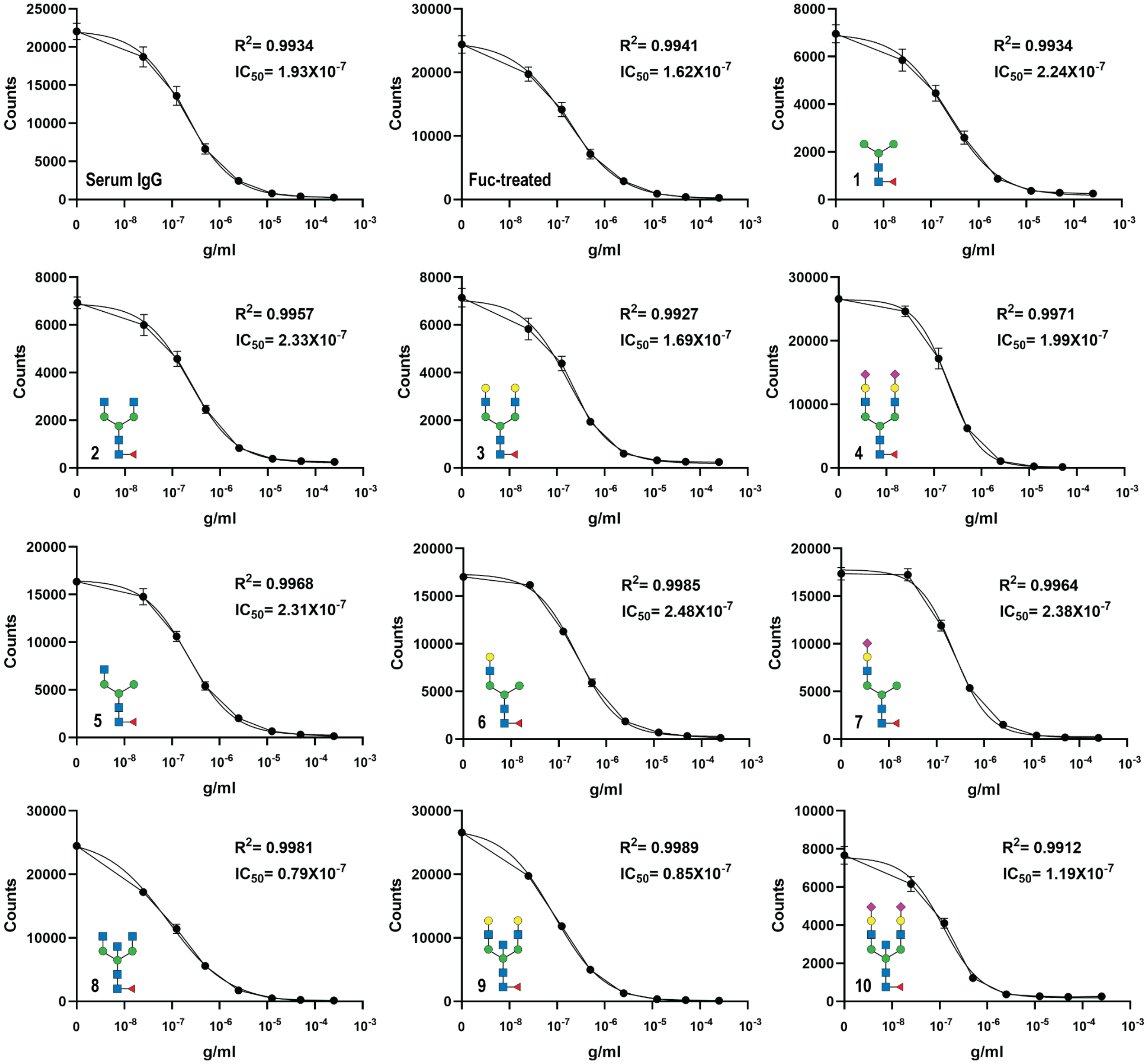

**Figure S13.** Investigating the binding affinity between SPGR-engineered IgG and Fcγ Receptor I using competition assays. IgG-FcRI interaction resulted in a signal decrease. Please note that all the IgG samples (**1**-**10**) contained a ~5% defucosylated population. The biantennary samples (**2**-**4**) had ~10% bisecting glycoforms; while the mono-antennary samples (**5**-**7**) also had ~10% (F)M3 glycans due to the GlcNAcase activity of GnT-I. Sialylated samples (**4** & **10**) possessed about 1:1 mono- and bi-sialylated populations. The Fuc-treated sample contained a 30% defucosylated glycan population. Effective concentration that leads to 50% signal reduction ($EC_{50}$) was calculated using $IC_{50}$ curve fitting by GraphPad Prism 8 (Dose-response, inhibition).

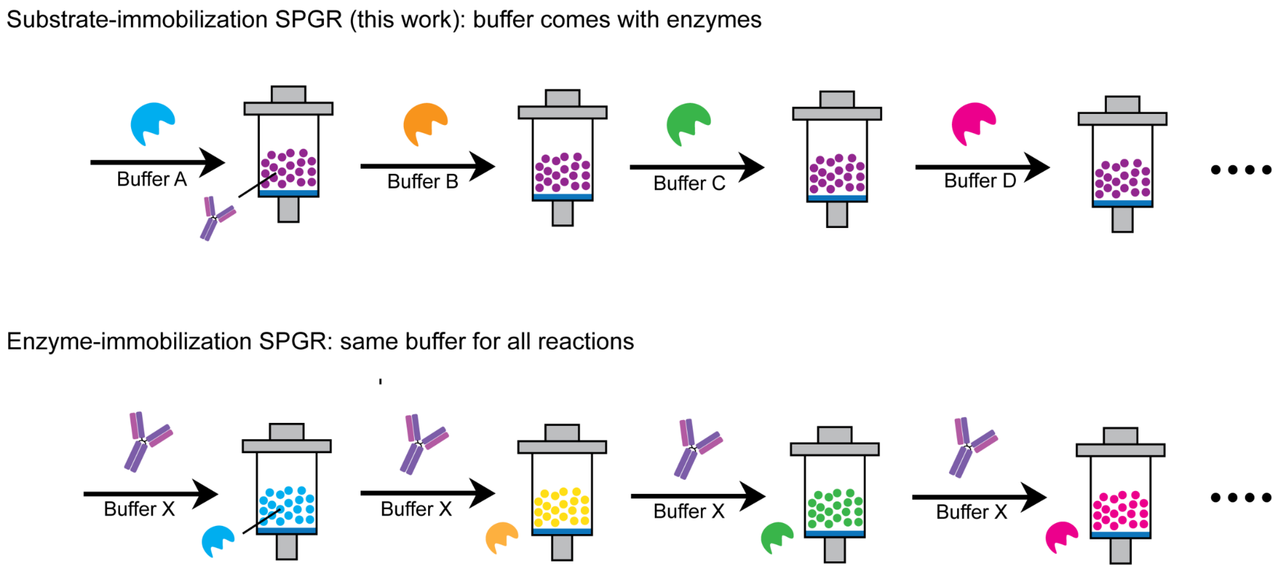

**Figure S14.** Comparison between substrate immobilization and enzyme immobilization in SPGR.

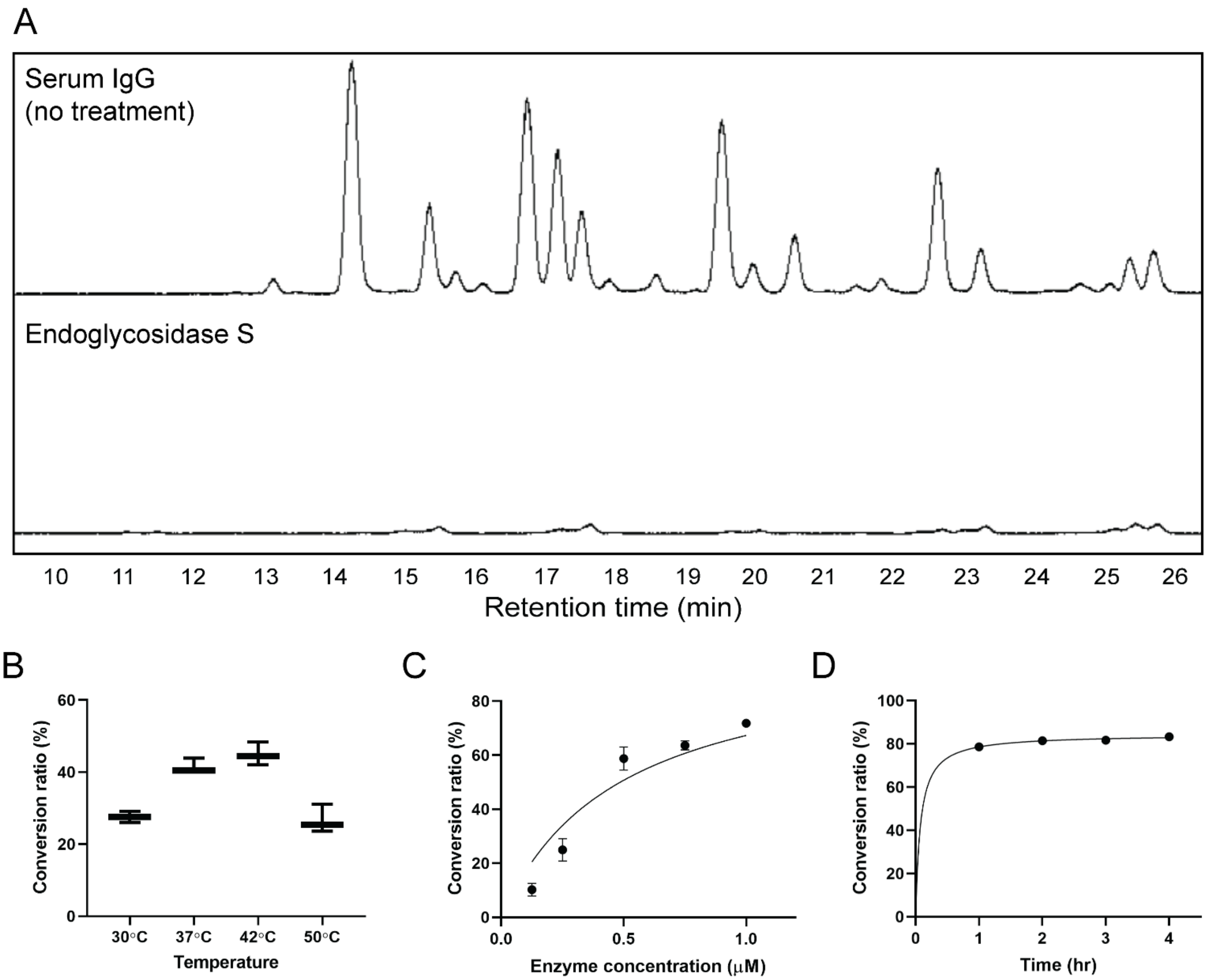

**Figure S15.** Characterization of endoglycosidase (*Streptococcus pyogenes*) activity and its working condition optimization. (A) Chromatography of IgG glycans before and after endo S treatment. (B) Temperature optimization. (C) Dose-dependent experiment. (D) Time-course study. Error bars: mean, maximum and minimum values. Refer to **Table S2** for reaction conditions.

Proprietary

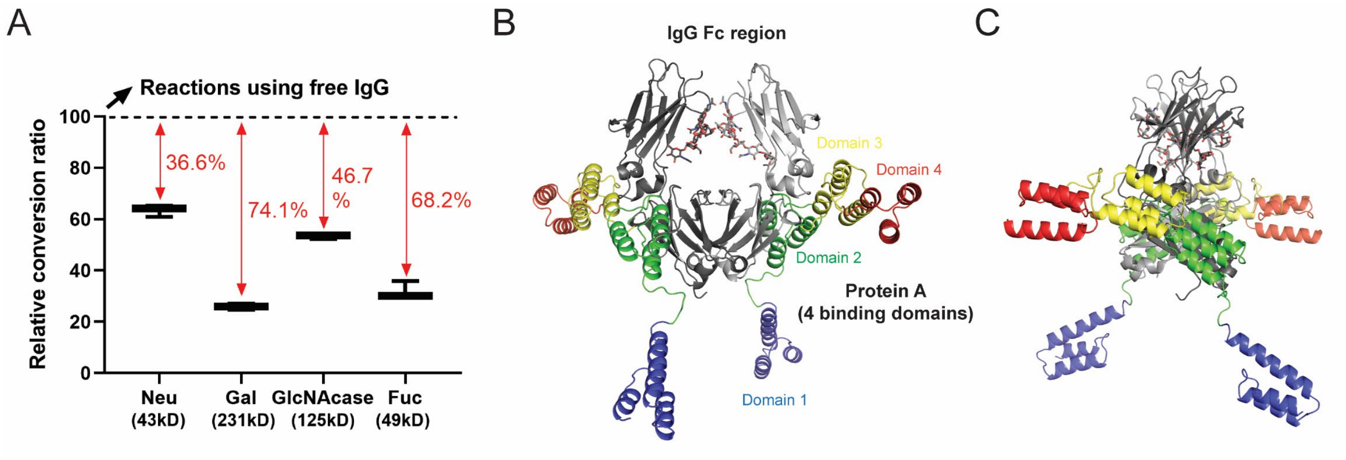

**Figure S16.** (A) Comparison of enzyme activity between the use of free and immobilized IgG as substrates. (B-C) Computational modeling of IgG Fc region and protein A interaction. (B) Front-view and (C) side-view of IgG Fc-protein A complex. Gray: IgG Fc region; blue: protein A binding domain 1; green: protein A binding domain 2; yellow: protein A binding domain 3; red: protein A binding domain 4. Bone structure: glycans.

### SI-References
